# Stabilized D_2_R G-protein coupled receptor oligomers identify multi-state β-arrestin complexes

**DOI:** 10.1101/2024.07.16.603735

**Authors:** Katie L. Sharrocks, Francesca Fanelli, Yewei Lui, Annabelle J. Milner, Wu Yining, Bernadette Byrne, Aylin C. Hanyaloglu

## Abstract

The G-protein coupled receptor (GPCR) superfamily directs central roles in many physiological and pathophysiological processes via diverse and complex mechanisms. GPCRs can exhibit signal pleiotropy via formation of di/oligomers both with themselves and other GPCRs. A deeper understanding of the molecular basis and functional role of oligomerization would facilitate rational design of activity-selective ligands. A structural model of the D2 dopamine receptor (D_2_R) homomer identified distinct combinations of substitutions likely to stabilize protomer interactions. Molecular modelling of β-arrestin-2 (βarr2) bound to predicted dimer models suggests a 2:2 receptor:βarr2 stoichiometry, with the dimer favouring βarr2 over Gαi coupling. A combination of biochemical, biophysical and super-resolution, single molecule imaging approaches demonstrated that the D_2_R mutant homomers exhibited greater stability. The mutant D_2_R homomers also exhibited bias towards recruitment of the GPCR adaptor protein βarr2 with either faster or ligand-independent βarr2 recruitment, increased internalization and reprogrammed regulation of ERK signaling. Through GPCR dimer-stabilization, we propose that D_2_R di/oligomerization has a role in βarr2-biased signaling.

## INTRODUCTION

G-protein coupled receptors (GPCRs) are the largest family of proteins in mammals, expressed in a range of cell types, and regulate a diverse set of physiological processes^1^. They are the targets of approximately 35% of current FDA approved drugs and remain major drug discovery targets^2,3^. GPCRs can regulate several intracellular signaling cascades through coupling with distinct heterotrimeric G-proteins. In addition, GPCRs can diversify their signaling through the multi-functional adaptor proteins, the non-visual arrestins; β-arrestin-1 and 2 (βarr1 and βarr2, respectively, also known as arrestins 2 and 3)^4,5^.In addition to uncoupling GPCRs from their heterotrimeric G-proteins, and driving receptor internalization via clathrin-coated pits, βarr1/2 can mediate distinct cellular pathways through activation of signaling such as the mitogen-activated protein kinase (MAPK) pathway ^4,5^.

Class A ‘rhodopsin-like’ GPCRs can function as monomers, however increasing biophysical, computational, pharmacological and physiological evidence suggests that they can form both homomers and heteromers with other GPCRs^6–12^. A dynamic equilibrium between monomers and transient di/oligomers has been reported for several GPCRs^13–16^. Oligomerization can impact the function of receptors, altering ligand efficacy, G-protein activation or preference for G-protein coupling selectivity^17–20^. The impact of oligomerization on GPCR/β-arr function is less clear but a recent study demonstrated that oligomerization of platelet-activating factor receptor (PAFR) decreases βarr recruitment and conversely, βarr seemed to decrease oligomerization of the receptor^21^. β_2_AR has also been shown to dimerise in a βarr-dependent manner upon stimulation with a βarr-biased ligand^22^. This highlights the potential for oligomerization to dramatically influence all aspects of GPCR signaling.

Structural biology remains an important area for uncovering GPCR activation and dimerization mechanisms^23^, yet there have been limited reports of high-resolution structures of Class-A GPCR dimers/oligomers. Transmembrane domains often play a key role in dimer interfaces, with the most recurrent interfaces involving helix 1 (H1). Indeed, H1-H1 contacts were found in homo-dimeric rhodopsin (PDB: 2I36, 2I37, 3CAP, and 6OFJ), κ-opioid (PDB: 4DJH), β1-adrenergic (PDB: 4GPO), M3-muscarinic (PDB: 5ZHP), EP3-prostanoid (PDB: 6AK3), AT1-angiotensin (PDB: 6DO1), and apelin (PDBs 6WOL and 6WON) receptors. Furthermore, H1-H4 contacts were found in homo-dimeric 5HT2A-serotonin (PDBs: 6WGT and 6WH4) and CCR5-chemokine (PDBs: 4MBS and 6AKX) receptors, whereas H1-H5 contacts were found in homo-dimeric D4 dopamine receptor (PDB: 6IQL). Homodimers characterized by H4-H5 or H4-H6 H5-H5, or H6-H7 were found for CXCR4, β1-AR, C5a1, CCR2, α2C-AR and CysLT1 receptors (PDBs: 3ODU/3OE8/3OE9, 5F8U, 5O9H, 6GPX, 6KUW, 6RZ5, respectively).

The D_2_ dopamine (D_2_R) is a class A ‘rhodopsin-like’ GPCR, coupling to Gα_i/o_^24^. The primary physiological roles of D_2_R in the brain are regulation of locomotion, and in reward and reinforcement mechanisms^25^, while pathophsyiologically this receptor is implicated in a number of neurological diseases including schizophrenia and Parkinson’s Disease^26^. D_2_R has been shown to form homodimers both in heterologous expression systems at physiological expression levels and in native tissues^14,27–29^. Several antipsychotics target the D_2_R/ βarr2 complex and an increase in D_2_R homodimers has been associated with schizophrenia^30,31^. Thus, D_2_R complexes represent excellent therapeutic targets. While there are several high-resolution structures of monomeric D_2_R, also in complex with G-protein^32–34^, the lack of information on precise molecular interactions between protomers and the absence of high-resolution dimer structures has hindered the ability to specifically target these complexes for therapeutic benefits.

Here we employed a convergent strategy to engineer distinct stabilized D_2_R homodimers. Residues were identified within the H1-H2 interface to enhance homodimer/oligomer stability assessed biochemically, biophysically and via super-resolution single molecule approaches. Interestingly, these engineered receptors also exhibited a bias towards βarr2 associated pathways with either altered temporal kinetics of ligand-mediated complexes, or functionally biased ligand-independent GPCR-arrestin complexes. Docking of βarr2 to the predicted homodimers of wild type (WT) or mutated D_2_R provide insights into the signaling active supramolecular assembly. This may highlight a novel selective role for D_2_R homomers in promoting distinct transitions to βarr2 association and signaling activity.

## RESULTS

### D_2_R mutations at the H2-H2 interface increase D_2_R homodimer stability and protomer proximity

To stabilize a D_2_R homodimer, our experimental design was guided by molecular modelling (see Methods). Simulations of D_2_R homodimerization were carried out using the structural models of either the inactive or the active states of the receptor. The results shown here concern symmetrical D_2_R homodimers with both protomers in their active states (i.e. extracted from the 8IRS PDB structural complex between ritigotine-bound D_2_R and heterotrimeric Gαi)), since they were most reliable in terms of membrane-topology indices. The predicted homodimer forms a symmetrical interface characterized by contacts at the extracellular ends of H1-H1, H1-H2 and H2-H2 and H1-H8, H8-H8 contacts on the cytosolic side (hereafter referred to as the H1, H2 dimer) (Fig. 1a). The stronger contacts from both protomers in the extracellular half of the interface involve: a) Y36 (H1) and F102 (extracellular loop 1 (EL1)); b) T39 (H1) and W90 (H2) and; c) T42 (H1) and I45 (H1) d) V97 (H2) from both protomers; and e) V96 and Y93 (both in H2). The same interface was found for symmetric D_2_R homodimers with both protomers in the inactive state (i.e. extracted from the 6CM4 PDB structural complex of D_2_R bound to risperidone) but not for asymmetric inactive-active D_2_R dimers. The presence of at least one protomer in the inactive state produced an additional interface characterized, among others, by H4-H4 contacts. For the H1,H2 dimer, we focused on the most extracellular amino acid residues to potentially drive formation of disulphide bridges. To strengthen the contacts between protomers at the interface, our models suggested mutations of Y93 to cysteine (Y93C), V96 to either serine (V96S) or cysteine (V96C) and V97 to cysteine (V97C) alone or in combination with V96S (V96S/V97C).

**Fig. 1.**
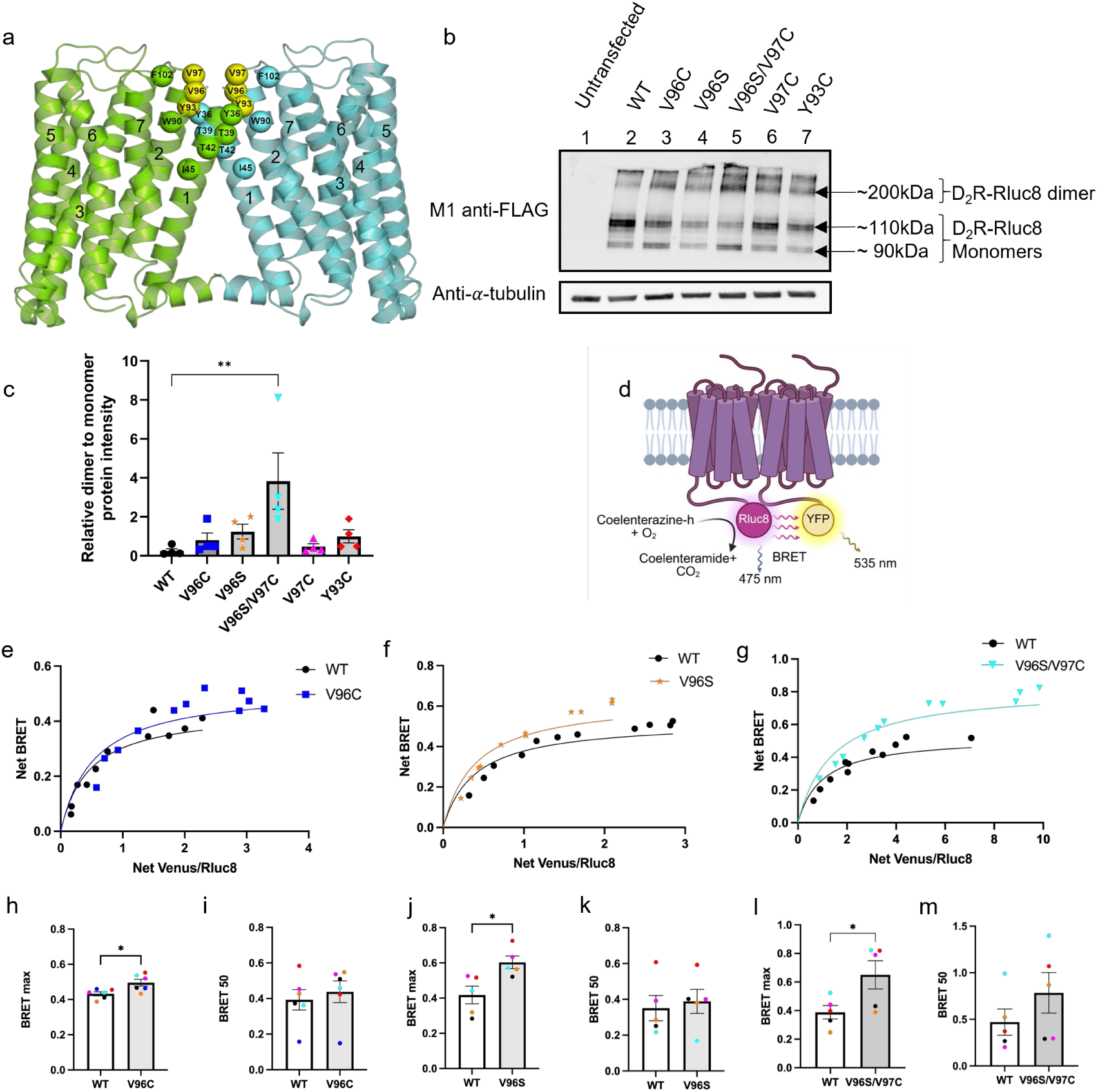
The impact of D_2_R mutants identified from molecular docking simulations on homodimer stability. **(a)** The predicted WT D_2_R homodimer (H1,H2 dimer) is shown in cartoon representation, with the two protomers coloured lemon-green and aquamarine. Interface amino acids are shown as spheres centred on the Cα-atoms. The mutated amino acids are highlighted in yellow. **(b)** Western blot analysis of HEK293 cell lysates transfected with WT or mutant D_2_R-Rluc8 as stated, carried out under reducing conditions. Probed with M1 anti-FLAG to detect the FLAG tag at the N-terminus of receptor constructs (D_2_R-Rluc8 monomer ∼90kDa or ∼110kDa with glycosylation) and anti-α-tubulin as a loading control. (**c**) Quantification of band intensity, represented as relative fold change of D_2_R dimer levels (∼200 kDa band) to monomer levels (∼110 kDa) under reducing conditions, N=4, error bars are +/-SEM. Ordinary one-way ANOVA used to measure statistical differences, (p**=0.0051) (**d**) Schematic showing BRET assay. (**e**) Representative BRET saturation curve obtained from HEK293 cells expressing either D_2_R WT (black), D_2_R V96C (blue), (**f**) V96S (orange) or **(g)** V96S/V97C (cyan). Saturation curves were used to quantity BRETmax (**h, j, l**) and BRET50 values (**i, k, m**), colour coded for each individual experiment. N=5 for WT vs V96S and V96S/V97C or N=6 for WT vs V96C. Un-paired, two-tailed Student’s t test used to measure statistical differences in BRET50 or BRETmax values between WT and mutant D_2_R, (p*= 0.0186 (h) or 0.0162 (j) or 0.0432 (l)).

To assess the effect of these mutations on levels of di/oligomeric protein present in cells, Western blot analysis under reducing or non-reducing conditions (in the presence or absence of β-mercaptoethanol) was carried out. Under non-reducing conditions receptors were primarily in a higher order, dimeric/oligomeric, complex (>180 kDa) (Supplementary Fig. 1). Under reducing conditions, the majority of the WT D_2_R shifted in size predominately to a monomeric form (predicted size of FLAG-tagged D_2_R-Rluc8, ∼90kDa with glycosylated D_2_R-Rluc8 appearing at ∼110kDa), however, a greater proportion of V96C, V96S and in particular, V96S/V97C mutants, existed in a higher order complex (Fig. 1b and c). Given the inability to distinguish between different types of higher-order oligomeric complexes using SDS-PAGE, quantification of only relative WT and mutant D_2_R dimer proteins levels of were compared (Fig. 1b and c). To directly measure receptor-receptor interactions in intact cells, bioluminescence resonance energy transfer (BRET) saturation assays were carried out. The WT D_2_R exhibited a saturation profile indicating formation of homomers (Fig 1e-g and Supplementary Fig. 2a and 3a), as previously reported by BRET^28^. When compared to WT D_2_R, a significant increase in BRETmax in cells expressing D_2_R V96C, V96S or V96S/V97C (Figs. 1e-m), indicating an increase in protomer proximity within D_2_R homomers. No significant changes in the receptor interaction profile were observed for Y93C (Supplementary Fig. 2b-d). All receptors exhibited a similar total expression level (Supp Fig. 2e-h) This supports the data obtained by Western blot analysis that the V96S/V97C mutation in particular may stabilize and/or increase the proportion of D_2_R homomers. There was no change in BRET50 (an indicator of protomer-protomer affinity) relative to WT for any of the mutants. Treatment of cells with the D_2_R-selective agonist, quinpirole, did not significantly impact either the BRETmax or BRET50 of WT or mutant D_2_Rs (Supplementary Fig. 3). To further understand how these sites impact receptor homomer interactions, computational simulations of V96C (Mut1) and V96S/V97C (Mut2) homodimerization were carried out and predicted that these mutated homodimers exhibit the same architecture as WT, though in Mut2 the interface is tighter than in WT or Mut1 (Supplementary Fig. 4). In the predicted Mut1 dimer, C96 interacts with both Y93 and C96 on the facing protomer, whereas in the Mut2 dimer C97 interacts with C97 and S96 interacts with Y93 on the facing protomer.

**Fig. 2.**
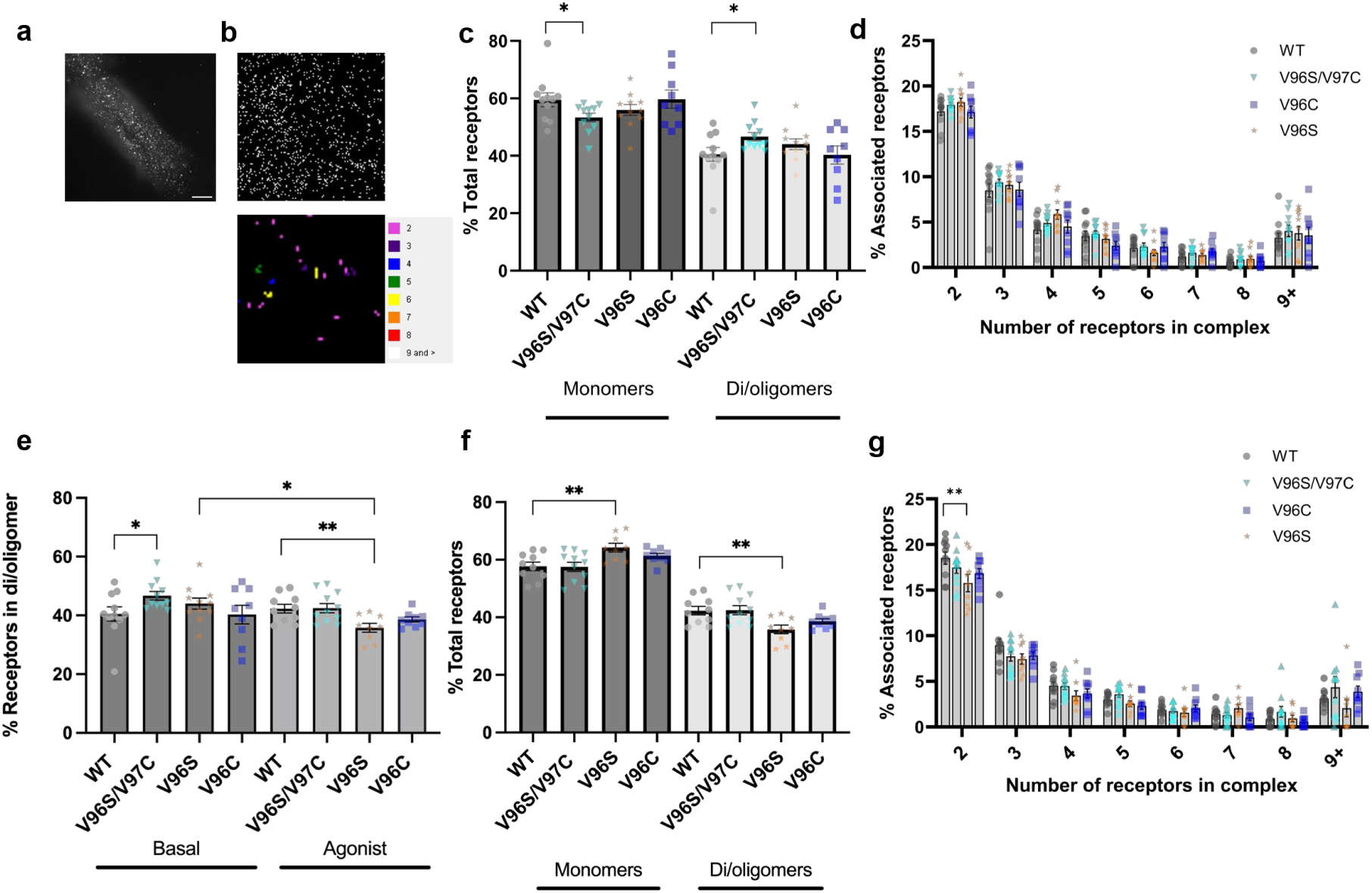
PD-PALM super-resolution microscopy allows quantification of cell surface D_2_R oligomers. (**a**) Representative image of HEK293 cells transiently transfected with FLAG-D_2_R and labelled with FLAG-CAGE500, imaged over 20,000 frames by TIRF microscopy following uncaging and activation of the CAGE500 dye. Scale bar=50 μm. **b)** Representative PD-PALM 7 μm^2^ image of FLAG-D_2_R labelled with FLAG-CAGE500 with a heat map quantifying oligomer populations. (**c**) Proportion of D_2_R WT or mutant receptors as monomers or in di/oligomeric complexes as a percentage of total receptors. (**d**) Composition of D_2_R oligomers separated into number of receptors in the complex and expressed as a percentage of total receptors. (**e**) Proportion of associated receptors in di/oligomeric complexes, expressed as a percentage of total number of receptors under basal conditions or after 5 minutes 10 μM quinpirole stimulation. (**f**) Proportion of D_2_R WT or mutant receptors as monomers or in di/oligomeric complexes as a percentage of total receptors after 5 minutes 10 μM quinpirole stimulation. (**g)** Composition of D_2_R oligomers after 5 minutes 10 μM quinpirole stimulation prior to fixation, separated into number of receptors in the complex and expressed as a percentage of total receptors. Mean +/-SEM, N=3 individual experiments, 3-4 cells imaged for each independent repeat. In c, e and f unpaired, two-tailed Student’s t test used to measure differences between WT and mutant D_2_R (p*=0.042 (c and e) or 0.0035 p**=0.0068 (e and f)). In d and g two-way ANOVA followed by Šídák’s multiple comparisons test used to measure statistical differences between WT and mutant D_2_R in different receptor complexes (p**=0.0019).

### Super-resolution imaging of cell surface D_2_R at the single molecule level reveals increased oligomerization in D_2_R V96S/V97C mutants

Both BRET and Western blot methods assess the total receptor population within a cell, whether in the biosynthetic, plasma membrane, or endocytic compartments. To quantitate the organisation of individual receptors at the plasma membrane, photoactivated localization microscopy with photoactivatable dyes (PD-PALM) with TIRF-imaging was employed. This single-molecule high-resolution technique gives <10 nm resolution and has provided detailed information on plasma membrane localised GPCRs in monomeric, dimeric or oligomeric forms^35–39^. PD-PALM imaging was carried out on HEK 293 cells expressing WT or mutant D_2_Rs and directly labelled with an anti-FLAG antibody conjugated to CAGE 500 at a 1:1 ratio. Precise localisation of individual receptors and subsequent processing through near neighbourhood analysis enabled individual monomer or oligomer populations to be quantified (Fig. 2a and b). Under basal conditions, the percentage of total receptors in a monomeric state in D_2_R V96S/V97C cells was significantly lower than WT D_2_R (Fig. 2c). Additionally, for V96S/V97C there was a corresponding significant increase in the proportion of receptors in the di/oligomeric state compared to WT D_2_R. When these di/oligomer populations were assessed according to the number of receptors in the complex, this revealed that these assemblies primarily involved receptor dimers (∼17% of total receptors), yet also a range of higher-order receptor oligomers. The increase in associated V96S/V97C receptor was not attributed to enrichment of a specific oligomer population but rather across receptor di/oligomers (Fig. 2d).

Following quinpirole stimulation, the different receptors exhibited distinct levels of monomer and oligomeric complexes, compared to basal conditions (Fig. 2e-g).

Stimulation of D_2_R WT expressing cells with quinpirole did not significantly alter the proportion of receptors in an oligomeric complex. However, ligand activation of D_2_R V96S resulted in a significant decrease in the proportion of receptor in an oligomer compared to both WT D_2_R expressing cells whose complexes were not significantly altered when stimulated with ligand (Fig. 2e and f). For V96S, this agonist-dependent decrease in associated receptors was due to a decrease in receptor dimers (Fig. 2g). While PD-PALM analysis suggests that quinpirole treatment may alter di/oligomerization of certain D_2_R mutants, quinpirole stimulation did not alter BRET saturation profiles (Supplementary Fig. 3 and Supplementary Fig.5). This could reflect that PD-PALM imaging quantifies complexes of cell surface receptors, while BRET detects interactions across all subcellular compartments.

We have previously demonstrated that receptor density can influence oligomerization for Class A GPCRs as measured by PD-PALM, particularly higher order complexes (>5 receptors/oligomer^39^). For D_2_R, the percentage of monomeric receptors was inversely proportional to the receptor density (Supplementary Fig. 5a and b). While for some GPCRs, receptor density impacts higher order complexes^39^, our observation for D_2_Rs via super-resolution imaging is in line with other studies with D_2_R_L_ using diffraction-limited single molecule imaging, reporting a small but significant impact of receptor density on the levels of monomer and di/oligomer distribution^14^. Therefore, based on the correlation observed via PD-PALM analysis, the di/oligomeric state of receptors were also analysed under ‘high’ (>100 receptors/μm^2^) or ‘low’ (<100 receptors/μm^2^) levels of expressed surface receptors, to understand the effect of receptor density on homomer formations in WT and mutant receptors (Supplementary Fig. 5b-h). In the absence of agonist, differences in complex formation between WT and mutant receptors were independent of receptor density. The range of receptor densities in cells treated with quinpirole was generally lower for cells expressing D_2_R V96S and V96C compared to WT or V96S/V97C, likely due to ligand-induced receptor internalization (Supplementary Fig. 5b). This may partially account for the decrease in oligomeric receptors in D_2_R V96S or V96C compared to WT D_2_R as when cells with similar receptor densities were analysed, this decrease was not significant (Supplementary Fig. 5g and h).

Taken together, this may suggest that the V96S/V97C mutant increases the propensity for D_2_R to form an oligomer. This is in line with docking simulations that predict a tighter interface in the V96S/V97C dimer compared to the WT or the V96C dimer (Supplementary Fig. 4). These results provide further evidence that the V96S/V97C mutant D_2_R may increase the stability of D_2_R oligomers. While quinpirole stimulation does not significantly alter the organization of D_2_R complexes.

### D_2_R mutants that stabilize homomeric interactions impact Gαi-mediated signaling and βarr2 recruitment profiles

Following the observation that some mutations increase the stability and/or degree of D_2_R homomers, the impact of these mutations on canonical Gαi signaling was assessed. Forskolin-induced cAMP levels were measured in the presence of quinpirole to assess inhibition of intracellular cAMP levels via D_2_R-mediated Gαi signaling. HEK293 cells expressing D_2_R WT and stimulated with 1 μM quinpirole for 5, 15, or 30 minutes exhibited a decrease in forskolin-induced cAMP levels at all agonist time points (Supplementary Fig. 6). This decrease in cAMP is specifically due to D_2_R-mediated signaling as untransfected HEK293 cells do not exhibit this decrease upon quinpirole addition. Concentration-response curves in cells expressing WT or mutant D_2_R showed similar curves and EC_50_ values, suggesting that the potency of quinpirole to activate D_2_R/Gαi-mediated signaling pathways is not altered by these mutations (Fig. 3a and Table 1). However, when the ligand-induced Gαi response was calculated from the change in cAMP levels in cells treated with the minimum to the maximum dose of quinpirole, the D_2_R V96S/V97C mutant had significantly reduced ligand-induced Gαi response compared to WT (Fig. 3b) suggesting a decrease in efficacy for quinpirole in D_2_R V96S/V97C expressing cells.

**Fig. 3.**
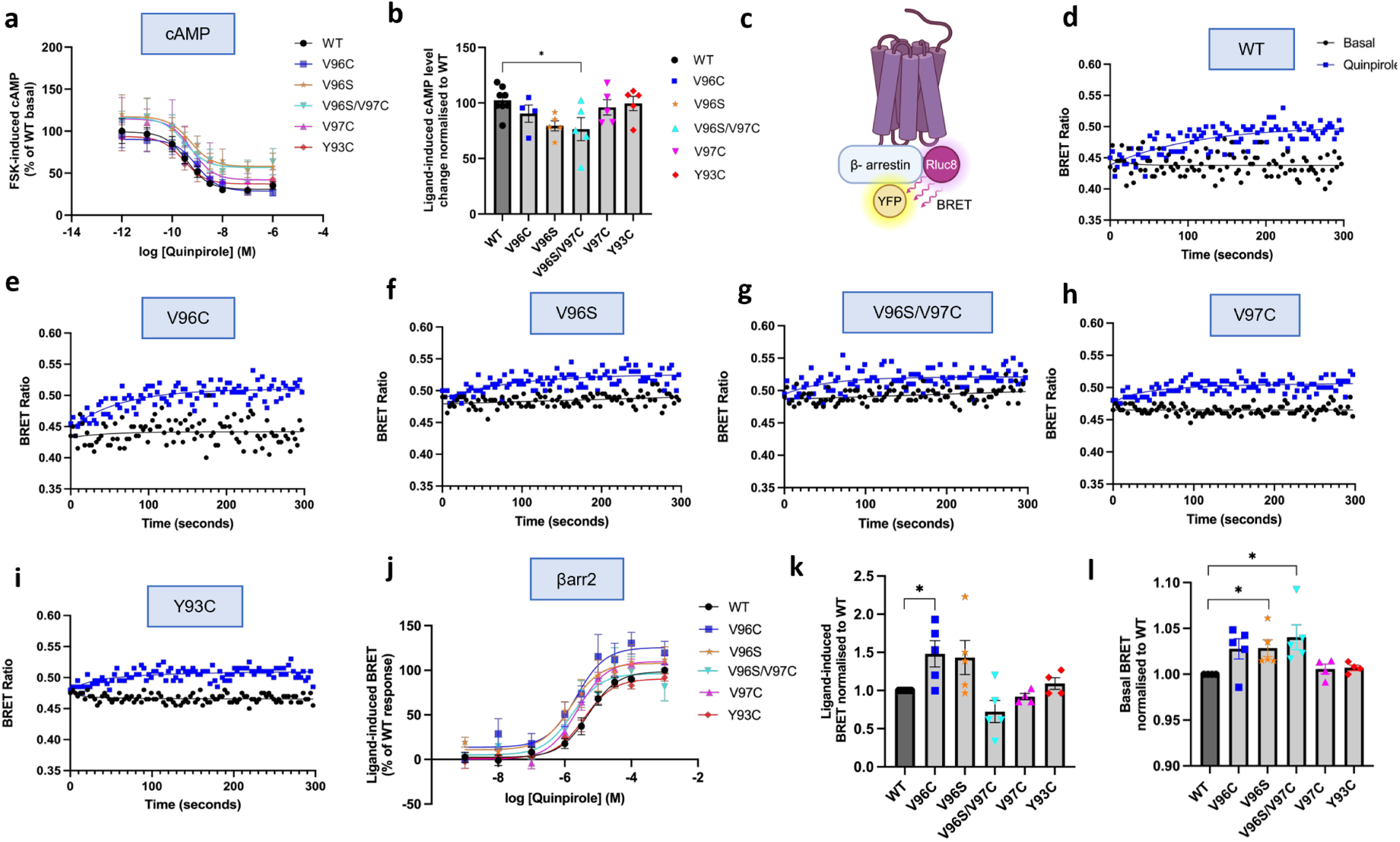
Altered G⍺i-signaling and βarr2 recruitment profiles of wildtype and mutant D_2_R. **(a)** Forskolin-induced cAMP levels as a percentage of basal cAMP levels of corresponding D_2_R WT, quinpirole concentration response curves in D_2_R WT (black) or mutant D_2_R transfected HEK293 cells. Values plotted are +/-SEM of 4-5 independent experiments for D_2_R mutants and 8 for WT. (**b**) Plot of quinpirole-induced cAMP levels, calculated as a change in cAMP levels in from cells treated with the maximum to the minimum dose of quinpirole, represented as a percentage of WT. One-way ANOVA and Dunnett’s multiple comparisons test used to measure statistical differences between WT and mutant maximal responses (p*= 0.00282 (V96S/V97C)) (**c**) Schematic showing BRET assay. Representative kinetic curve showing BRET ratios in HEK293 cells expressing βarr2-YFP and D_2_R WT (**d**), V96C (**e**), V96S (**f**), V96S/V97C (**g**), V97C (**h**), Y93C (**i**). Plotted over time for basal (black) and after 10 μM quinpirole stimulation (blue). Graphs representative of 4-6 independent experiments. (**j**) BRET luminescence values were measured before agonist addition and after addition of 1 nM – 1mM quinpirole, values are average of measurements taken over 5 minutes. Represented as a percentage of D_2_R WT maximal response, for each individual experiment. (**k**) To determine the efficacy of quinpirole to induce βarr2 recruitment, for each individual experiment, ligand-induced BRET values were plotted for cells treated with the maximum dose of quinpirole (1 mM), represented as a fold change from D_2_R WT. Basal BRET (**l**) of cells with no ligand and only PBS added prior to luminescence measurement for each individual experiment was plotted and presented as a fold change from WT. One sample t and Wilcoxon test. N=4 (V97C and Y93C) or 5 (V96C, V96S and V96S/V97C), +/-SEM. (p*= 0.0488 (k) or 0.0344 or 0.0398).

**Table 1.** Quinpirole potency in cAMP and βarr2 BRET recruitment assays shown as –logEC_50._ Data shown with 95% confidence interval (CI). Two-tailed, unpaired Student’s t-test used to determine statistical significance between WT and mutant –logEC50 values (p*=0.0147 (V96C) or 0.0433 (V96S)).

| Receptor | cAMP $-\log EC_{50}$ | βarr2 recruitment $-\log EC_{50}$ |
| --- | --- | --- |
| D<sub>2</sub>R WT | -9.418 (CI -9.761 to -9.073) | -5.362 (CI -5.527 to -5.196) |
| D<sub>2</sub>R V96C | -9.036 (CI -9.535 to -8.546) | -5.695 (CI -5.814 to -5.576)* |
| D<sub>2</sub>R V96S | -9.275 (CI -10.280 to -8.358) | -5.804 (CI -6.044 to -5.565)* |
| D<sub>2</sub>R V96S/V97C | -9.490 (-10.270 to -8.687) | -5.622 (-5.788 to -5.457) |
| D<sub>2</sub>R V97C | -9.225 (CI -10.020 to -8.357) | 5.626 (CI -5.167 to -6.0850) |
| D<sub>2</sub>R Y93C | -9.547 (-10.180 to -8.920) | 5.435 (CI 5.242 to 5.628) |

To assess the impact of these mutant D_2_Rs on βarr1/2 recruitment, BRET assays were carried out in HEK293 cells expressing D_2_R and βarr1/2. The kinetics of βarr1/2 recruitment were analysed in real time via BRET measured over a time-course of 5 minutes following agonist stimulation. The WT D_2_R did not recruit βarr1 following quinpirole treatment (Supplementary Fig. 7a and c). However, in cells expressing D_2_R WT and βarr2-YFP there was a rapid increase in BRET levels specifically after quinpirole addition, up to a maximum level of BRET signal which remained stable for at least 5 minutes (Fig. 3d and Supplementary Fig. 8). All D_2_R mutants (except for D_2_R-Rluc8 V96S/V97C) showed a similar profile (Fig. 3e-i). The ability of D_2_R to couple to βarr2 over βarr1 is consistent with other studies^40,41^. Calculation of ligand-induced BRET values and subsequent curve-fitting of the BRET profile allowed quantification of the kinetics of βarr2 recruitment to WT and mutant D_2_Rs (Table 2 and Supplementary Fig. 8). There was no significant difference in the plateau of the fitted kinetic profiles between WT or any mutant D_2_R. However, the halftime of the D_2_R V96C kinetic curve was significantly lower than WT, suggesting that D_2_R V96C exhibits altered βarr2 recruitment.

**Table 2.** Quinpirole-induced βarr2 BRET recruitment kinetics. Rate (k), initial rate, halftime and plateau were calculated following curve fitting from the waveform produced from ligand-induced BRET recruitment assays (Supplementary Fig.6). Data is presented with 3 different D2R WT data sets, to allow comparisons with mutant D2R experiments that were carried out at the same time under the same experimental conditions. N=4-6, data shown as +/-SEM. Two-tailed, unpaired Student’s t-test used to determine statistical significance between WT and mutant-logIC50 values (p**= 0.0014 or 0.0068).

|  | k (second<sup>-1</sup>) | Initial Rate (BRET ratio units per second) | Half time (seconds) | Plateau |
| --- | --- | --- | --- | --- |
| **WT** | $1.076 \times 10^{-2} (\pm 1.283 \times 10^{-3})$ | $5.27 \times 10^{-4} (\pm 1.644 \times 10^{-4})$ | 68.38 ($\pm 8.537$) | 0.04908 ($\pm 1.166 \times 10^{-2}$) |
| **V96C** | $1.999 \times 10^{-2} (\pm 1.446 \times 10^{-3})^{**}$ | $1.002 \times 10^{-3} (\pm 1.842 \times 10^{-4})$ | 35.55 ($\pm 3.058$)<sup>**</sup> | 0.05012 ($\pm 8.027 \times 10^{-3}$) |
| **WT** | $4.754 \times 10^{-3} (\pm 3.549 \times 10^{-3})$ | $1.579 \times 10^{-4} (\pm 1.049 \times 10^{-4})$ | 30.28($\pm 4.99$) | 0.03682 ($\pm 7.335 \times 10^{-3}$) |
| **Y93C** | $3.33 \times 10^{-3} (\pm 1.943 \times 10^{-3})$ | $1.135 \times 10^{-4} (\pm 5.866 \times 10^{-5})$ | 34.57($\pm 14.05$) | 0.03119 ($\pm 7.849 \times 10^{-3}$) |
| **V97C** | $4.434 \times 10^{-3} (\pm 3.577 \times 10^{-3})$ | $1.807 \times 10^{-4} (\pm 1.499 \times 10^{-4})$ | 32.66 ($\pm 4.764$) | 0.0297 ($\pm 6.017 \times 10^{-3}$) |
| **WT** | $3.065 \times 10^{-2} (\pm 1.072 \times 10^{-2})$ | $8.603 \times 10^{-4} (\pm 5.241 \times 10^{-4})$ | 41.08($\pm 7.252$) | 0.02122 ($\pm 5.301 \times 10^{-3}$) |
| **V96S** | $2.702 \times 10^{-2} (\pm 8.711 \times 10^{-3})$ | $7.002 \times 10^{-4} (\pm 3.117 \times 10^{-4})$ | 39.36 ($\pm 8.69$) | 0.02394 ($\pm 3.826 \times 10^{-3}$) |

To assess if the potency of quinpirole to induce βarr2 recruitment to mutant receptors was altered, cells were treated with increasing concentrations of quinpirole (Fig. 3j-l). D_2_R V96C and V96S receptors, but not V97C or Y93C, exhibited a leftward shift in the concentration-response curves, compared to WT with significantly lower EC_50_ values (Table 1). This suggests that in cells expressing the V96C and V96S mutants, quinpirole exhibits increased potency towards βarr2 recruitment. Additionally, there is a significant increase in basal βarr2 recruitment in cells transfected with D_2_R V96S and V96S/V97C compared to D_2_R WT (Fig. 3l). Combined with the reduced ligand-induced BRET signal in kinetic assays (Fig. 3), this may suggest increased constitutive association between the V96S/V97C receptor and βarr2. These signaling changes were not due to alterations in cell surface receptor expression as reducing D_2_R WT cell surface expression to levels obtained with the mutant receptors had no impact on the signal activity profile (Fig. 4 and Supplementary Fig. 9). Given that certain D_2_R mutants exhibit both stabilized/enhanced homomerization and a potential bias to βarr2, we then assessed whether UNC9994, a D_2_R-selective ligand, which has been reported to show biased βarr2 signaling in some cell contexts, impacted D_2_R WT signaling and homomerization (Supplementary Fig. 10)^42,43^. A significant decrease in D_2_R-mediated Gαi signaling was observed and while a more transient kinetic profile was observed for βarr2 recruitment, this was not enhanced compared to quinpirole, suggesting UNC9994 acts as a partial biased agonist in this context. Additionally, UNC9994 did not impact D_2_R homomer associations as indicated by the BRETmax or BRET50 values, similar to what was observed following quinpirole stimulation (Supplementary Fig. 10 e-g).

**Fig. 4.**
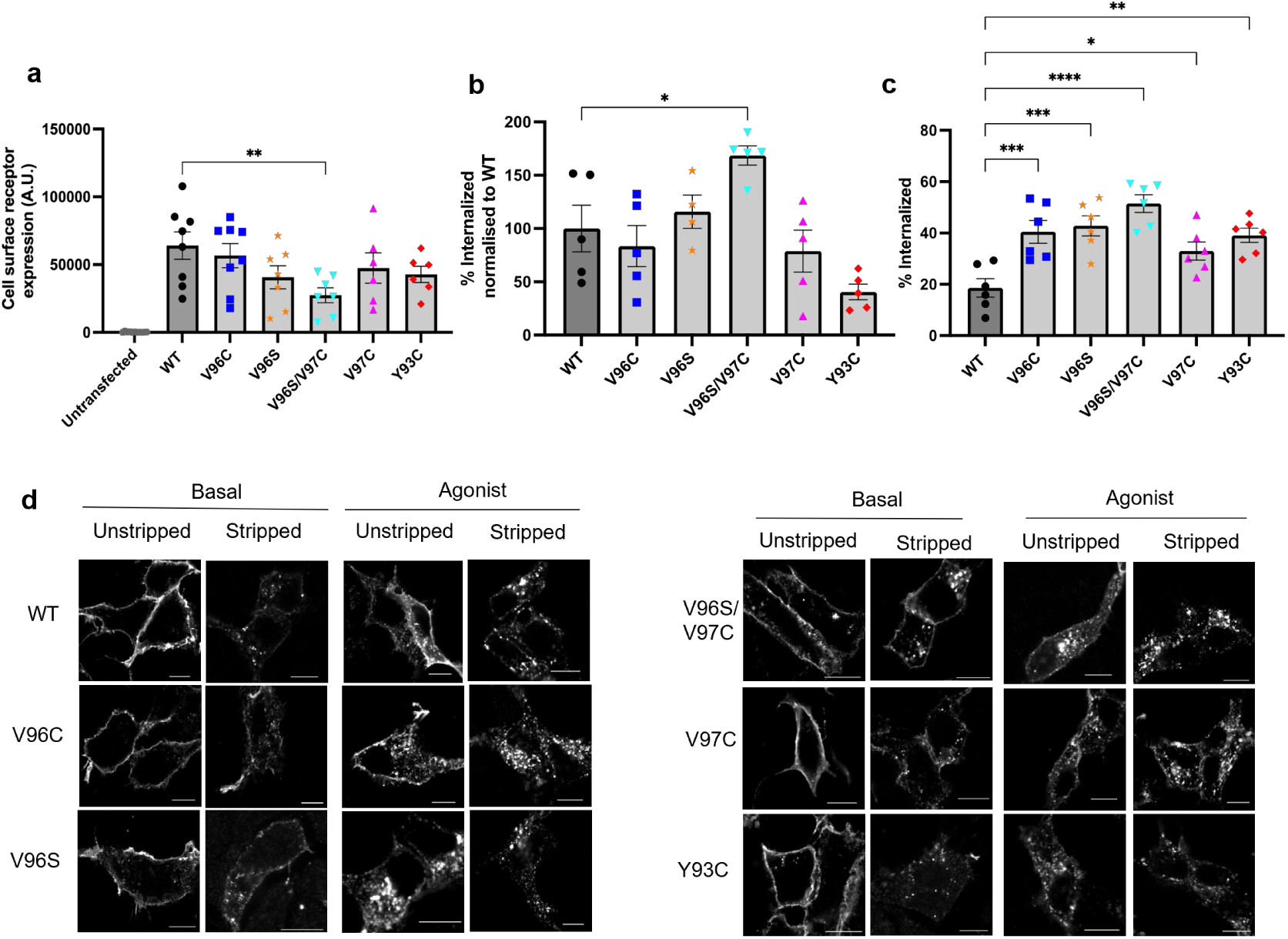
Receptor cell surface expression and internalization assessed by flow cytometry and immunofluorescence microscopy. HEK293 cells transfected with D_2_R constructs were labelled with M1 anti-FLAG primary and Alexa647 conjugated secondary antibody to quantify cell surface receptor expression. (**a**) Mean fluorescence and % of gated cells were multiplied to give a total value for cell surface D_2_R expression. N=6-8. (**b**) In cells expressing βarr2-YFP, constitutive internalization of receptor constructs was calculated as a percentage change of cell surface receptor expression in cells incubated at 4°C to prevent receptor internalization, compared to cells incubated at 37°C following antibody labelling, which can continue to constitutively internalize receptor. Expressed as a percentage of WT for each individual experiment. N= 4 or 5. (**c**) Quinpirole-induced receptor internalization. Measured as a percentage decrease in cell surface expression level following 10 μM quinpirole stimulation, compared to untreated cells. N=6. In a-c, error bars are +/-SEM. One-way ANOVA followed by Dunnett’s multiple comparison test used to measure statistical differences between WT and mutant D_2_R (p*=0.0273 (b) or 0.0369 (c), p**= 0.0065 (a) or 0.0018 (c), p***<0.0009 or 0.0003, p**** <0.0001. (**d**) Representative confocal microscopy images showing D_2_R localisation under basal conditions and after 30 minutes 10 μM quinpirole stimulation, with and without ‘stripping’ of cell surface bound anti-FLAG antibody. Representative of 3 independent experiments. Scale bar= 10 μm.

Overall, this data suggests that V96C, and to some extent V96S show altered quinpirole-induced βarr2 recruitment to D_2_R, while V96S/V97C exhibits enhanced constitutive association of βarr2. Furthermore, these receptors exhibit bias towards βarr2 recruitment over Gαi signaling pathways.

### Constitutive and ligand-induced receptor internalization is enhanced in D_2_ R V96S/V97C mutants

β-arrestins play key roles in regulating GPCR signal activity^44,45^. One function is driving receptor-mediated internalization. Given the faster (V96C) βarr2 recruitment and, in particular, the ligand-independent (V96S/V97C) recruitment of βarr2, the cell surface expression of these D_2_Rs was assessed via flow cytometry and confocal microscopy. While all receptors were shown to traffic to the membrane (Supplementary Fig. 11), the cell surface expression of V96S/V97C was lower than WT D_2_R (Fig. 4a). Due to the observation that V96S/V97C also shows increased basal βarr2 recruitment, the possibility that this decrease in cell surface expression was due to increased internalization was investigated. The percentage of constitutively internalized receptor was calculated as a percentage decrease of cell surface expression in cells incubated at 37 °C following live antibody labelling, compared to cells incubated at 4°C to prevent receptor internalization (Fig. 4b). Cells were gated to quantitate levels of D_2_R in βarr2-YFP-expressing cells. An increase in constitutive internalization of V96S/V97C compared to WT receptor was observed, while Y93C had decreased internalization, and all other mutants showed comparable levels of internalization to cells expressing WT receptor. This increase in constitutive internalization may at least partially account for the decrease in cell surface expression for D_2_R V96S/V97C compared to WT. There was a significant increase in ligand-induced internalization for all mutant receptors, compared to WT and this increase was most dramatic in D_2_R V96S/V97C-expressing cells (Fig. 4c).

The altered ligand-induced and constitutive internalization in mutant D_2_Rs observed quantitatively by flow cytometry, was also confirmed by confocal microscopy (Fig. 4d).

Given the decrease in cell surface expression of some D_2_R mutants, the impact of lower cell surface D_2_R expression on internalization and signaling was assessed. Transfection of lower amounts of WT D_2_R plasmid resulted in lower cell-surface receptor expression, however, this decrease in WT expression did not cause a significant change in ligand-induced or constitutive internalization or the efficacy of quinpirole-mediated Gαi signaling (Supplementary Fig. 9c-e), suggesting increased internalization and decreased Gαi signaling in receptor mutants is not solely due to lower cell surface expression. Additionally, reduction WT D_2_R expression to a comparable level to the V96S/V97C receptor, results in the opposite effect of a decrease in basal associations with βarr2, suggesting that the observed increase in basal BRET associations with βarr2 in V96S/V97C-expressing cells compared to WT is not solely due to lower expression levels of mutant receptor (Supplementary Fig. 9a and b and Fig. 3). This data suggests that enhanced basal βarr2 association may promote both ligand-independent and ligand-induced internalization.

### D_2_Rs that promote βarr2 recruitment favor βarr2-mediated ERK signaling

Arrestins play well-documented roles in mediating activation of extracellular signal-regulated protein kinase (ERK) signaling^46^, therefore, the impact of these mutations on the phosphorylation levels of ERK1/2 was assessed. D_2_Rs WT, V96S and V96S/V97C increased ERK activation up to a maximal level following 2-5 minute stimulation with quinpirole (Fig. 5a and b). However, V96C exhibited enhanced agonist-dependent ERK signaling compared to WT at the 5-minute quinpirole stimulation timepoint, while V96S/V97C exhibited a more rapid ERK signaling profile (Fig. 5a and b). In CRISPR βarr1/2 knockout-cells, the ability of WT D_2_R to increase phospho-ERK levels after quinpirole stimulation was maintained (Fig. 5c-e). D_2_R V96C, V96S and V96S/V97C, however, exhibited a decrease in agonist-dependent ERK signaling (Fig. 5f-h and Supplementary Fig. 12). This sensitivity/requirement of βarr for ERK signaling could be due to the altered βarr2 association profiles of these D_2_R mutants, which also in turn exhibit enhanced/stabilized homomer associations.

**Fig. 5.**
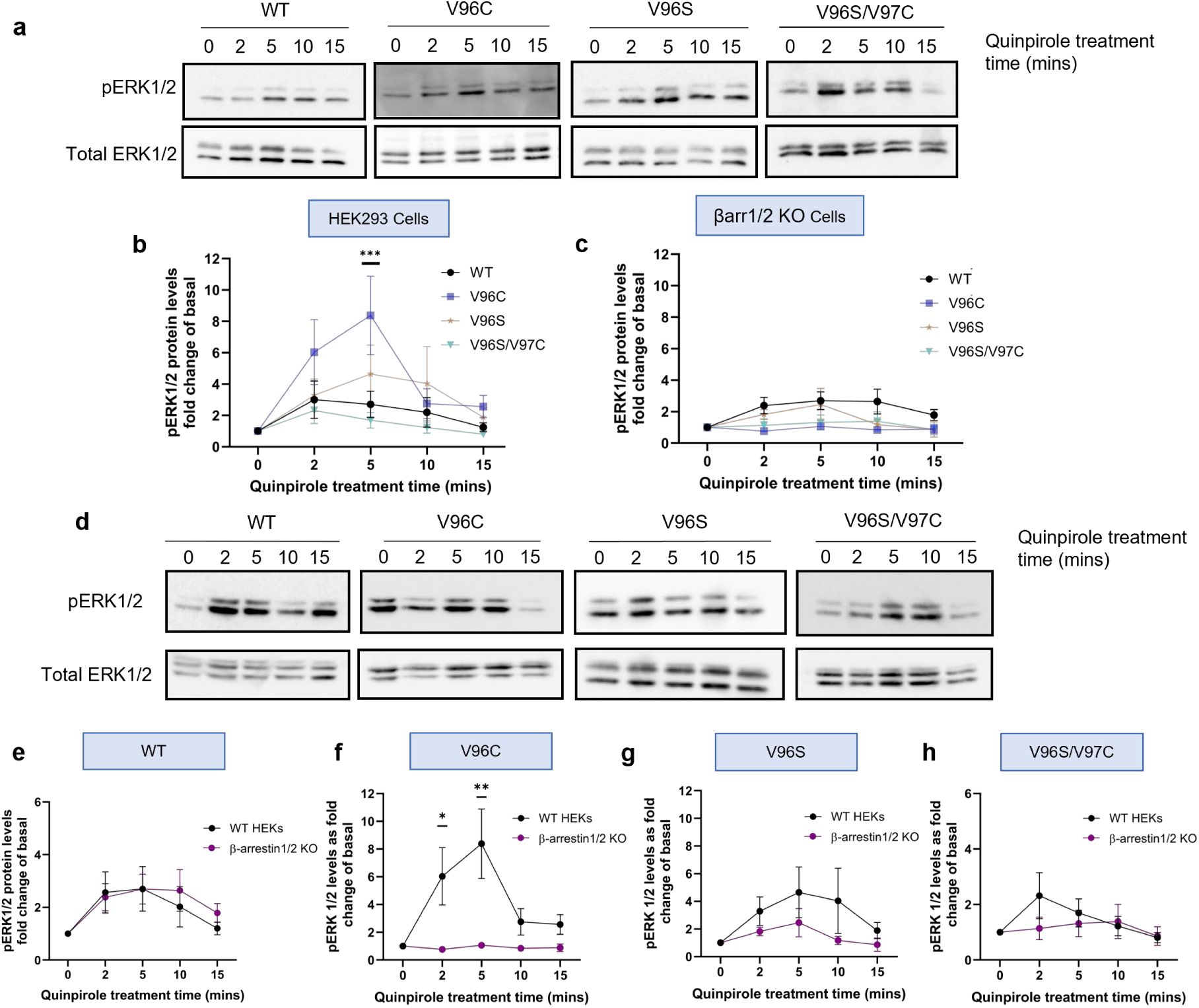
Ligand induced ERK1/2 activation in HEK293 cells and βarr 1/2 knockout cells. Representative Western blots on HEK293 cells (**a** and **b**) or HEK293 βarr 1/2 KO (βarr 1/2 KO) cell lysates (**c** and **d**) transfected with D_2_R constructs and treated with 10 μM quinpirole for 2, 5, 10 or 15 minutes, probed with phospho-ERK 1/2 and total ERK 1/2 antibodies. Normalised ligand-induced phospho-ERK 1/2 levels expressed as a fold change of basal levels quantified in HEK293 (**b**), or βarr 1/2 KO cell lysates (**c**) transfected with D_2_R-Rluc8 WT (black) vs V96C (blue), V96S (orange), V96S/V97C (turquoise) constructs. (**e-h**) Direct comparison of relative pERK1/2 levels in WT HEK293 (black) or βarr 1/2 KO cells (purple) transfected with D_2_R WT (**e**), V96C (**f**), V96S (**g**), V96S/V97C (**h**). Mean +/-SEM, N=5 (V96S and V96S/V97C) or 6 (V96C) for mutant receptors in HEK293 cells and 4 (V96C and V96S/V97C) or 5 (V96S) in βarr 1/2 KO cells and N=8 for WT D_2_R. Two-way ANOVA followed by Šídák’s multiple comparisons test used to measure statistical differences between WT and mutant D_2_R (in b p***<0.0005, in f p*= 0.0439 and p**=0.015).

### Differential stoichiometry of D_2_ R dimers with its G proteins or arrestin

Insights into the architecture of the assemblies between dimeric D_2_R and heterotrimeric GαI or βarr2 were inferred from molecular modelling. By fitting the D_2_R from the cryoEM structure in complex with heterotrimeric Gαi (PDB: 8IRS) onto both protomers in each of the predicted D_2_R homodimers, a 2:1 D_2_R:Gαi stoichiometry is predicted (Supplementary Fig. 13). Steric clashes occur between two G-protein heterotrimers simultaneously docked to the D_2_R homodimer, making a 2:2 stoichiometry unlikely. This suggests that for dimeric D_2_R the proper D_2_R:Gαi stoichiometry is 2:1 and any stabilization at the H1-H2 D_2_R homodimer, especially V96S/V97C, is expected to reduce the number of receptors available for binding to heterotrimeric Gαi, thus reducing coupling efficiency. The 2:1 D_2_R:Gαi stoichiometry is in line with the minimal signaling unit found to maximally be activated by agonist binding to a single protomer in an asymmetrical activated D_2_R dimer ^47^.

Docking simulations between the homodimers of D_2_R WT, V96C, and V96S/V97C and two different structural models of βarr2 always predicted receptor-arrestin complexes similar to the cryoEM complexes (see Methods). It is worth noting that in all dimeric complexes, the D_2_R is in an active state as it was extracted from the cryoEM complex with the agonist ritigotine and heterotrimeric Gi (PDB: 8IRS). Remarkably, the architecture of the H1-H2 D_2_R dimers favors the simultaneous recruitment of two βarr2 molecules (i.e. with a D_2_R:βarr2 2:2 stoichiometry) (Fig. 6a and b). Noticeably, the finger loop holds a different conformation in the two βarr2 models, which was inherited from the two different βarr1 structural templates, the finger loop being unstructured in 6TKO while holding one α-helical turn in 7SRS. In the predicted complexes between WT or mutant D_2_R dimers and both βarr2 structural models, the central-crest loops of active βarr2 form a docking site for IL2 of the receptor, which holds a two-turn α-helix. Specifically, the finger loop of βarr2 docks in the receptor site contributed by the cytosolic extensions of H3 and H6 as well as IL1, IL2, and H8. Furthermore, the middle loop and the β-hairpin that includes the C-loop of βarr2 make contacts with IL2 and H5 of the receptor (Fig. 6a and b). While the interface described above is common to all prototypical GPCRs, an additional interface between the N-terminal domain of βarr2 and the phosphorylated tail of the receptor is formed only when such tail is present, which is not the case of D_2_R.^48^ Lack of the phosphorylated C-tail may have an effect on the process of βarr-receptor recognition but not on the final architecture of the complex, which is expected to be the same independent of the presence or absence of the C-tail. Therefore, the D_2_R dimer-stabilizing mutants both reduce the number of receptors available for Gαi coupling, whilst favoring βarr2 coupling. Measurement of D_2_R homomer interactions via BRET saturation assays in βarr½2 KO cells suggested that the lack of βarr1/2 did not significantly alter D_2_R dimerization (Supplementary Fig. 14). This may suggest that D_2_R dimers form distinct conformations in the presence or absence of βarrs, without changing overall dimer stability.

**Fig. 6.**
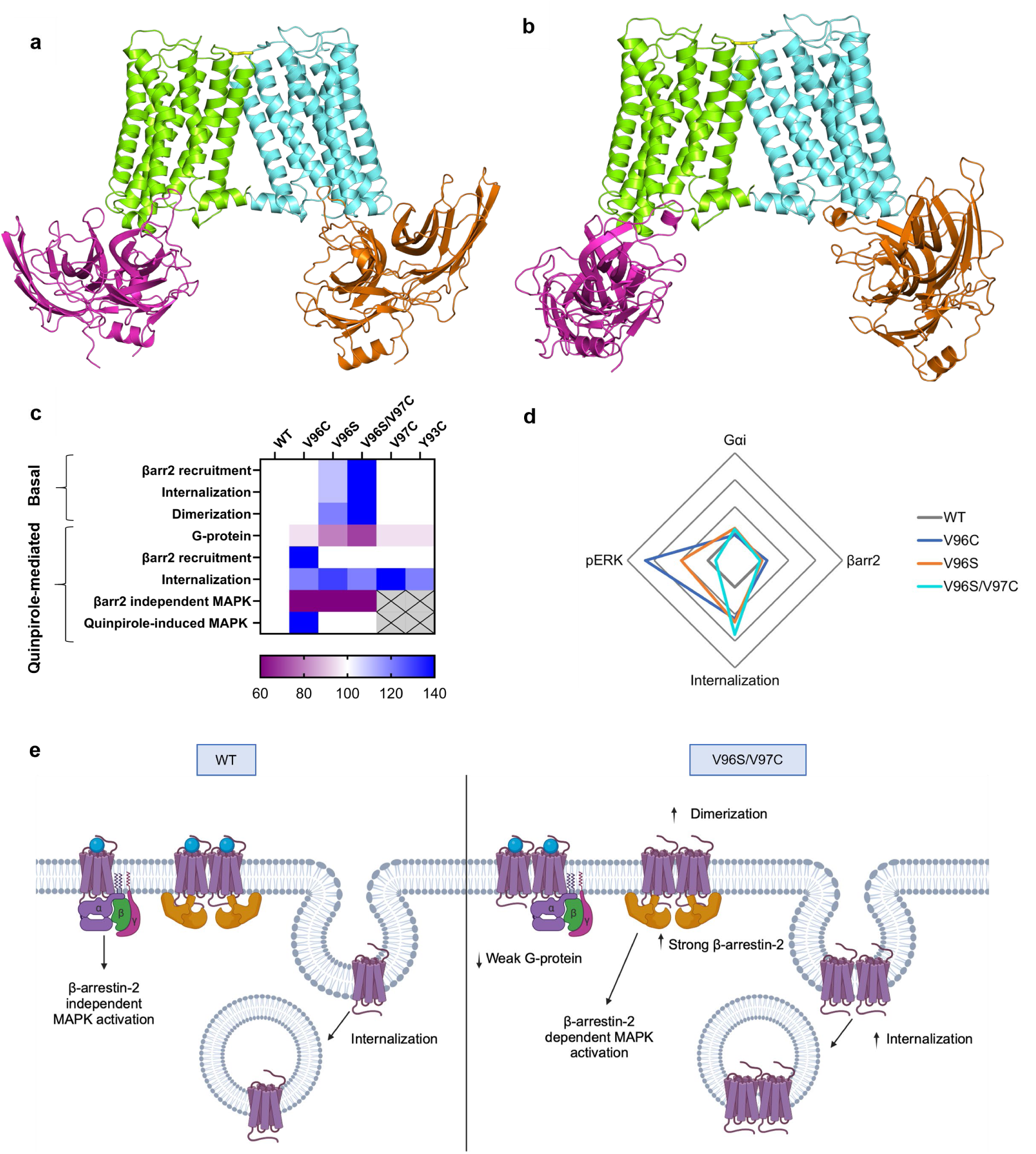
Predicted models for D_2_R homodimer-βarr2 complexes and summary schematics. The cartoon representations of two different structural complexes between the V96S/V97C mutant D_2_R homodimer (protomers lemon-green and aquamarine) and βarr2 ( orange and magenta) are shown. The structural models of βarr2 shown in the **a** and **b** panels have been, respectively, achieved by using the βarr1 structures from the 6TKO and 7SRS cryoEM complexes as templates. The side chains of C97 are shown in yellow sticks; the distance between the two sulfur atoms is 2.0 Å. I) Heat map summarising the functional effects of D_2_R mutants relative to WT. 100 is WT and white, an increase relative to WT is blue and a decrease is purple. (**d**) Quinpirole induced Gi signalling, βarr2 recruitment, quinpirole-induced internalization and pERK activation of D_2_R mutants as a fold change relative to D_2_R WT. G-protein and β-arrestin bias calculated as Emax/EC_50_ (**e**) Summary schematic of proposed βarr2 biased signalling from D_2_R V96S/V97C oligomers.

## DISCUSSION

Combining molecular docking simulations with biophysical, super-resolution and functional studies, we propose a model for stabilizing D_2_R homomer associations and demonstrate the utility of such an approach to identify a mechanism for not only stabilizing receptor-receptor associations, but also for promoting both βarr2 interaction and βarr2-mediated functions. This reveals a potential role for D_2_R di/oligomerization in βarr2-biased signaling that could be targeted for therapeutic benefits.

The predicted interface in symmetric D_2_R homodimers with both protomers either inactive or active, characterised by H1-H1, H1-H2, and H2-H2 contacts (i.e. H1,H2 dimer), is shared with other published high-resolution GPCR dimer structures including rhodopsin, κ-opioid, β1-adrenergic, μ-opioid, and human apelin receptors^49–53^. Previous studies have suggested that H4 participates in D_2_R homodimer interface, but a second interface involving H1 and H8 has also been suggested^28,54,55^. We found the involvement of H4 in some D_2_R dimers with both protomers inactive or with asymmetric inactive-active dimers, though with worse membrane topology. Additionally, several studies have suggested that IL1, EL2 and EL3 are important in stabilizing GPCR dimer interactions^55–57^. However, this is based on computational modelling and biophysical data, with no current published dimer structure for the D_2_R homodimer. It is also possible that multiple dimer interfaces exist, as reported for the β1AR, where both a H1-H2-H8-EL1 and H4-H5-EL2-IL2 interfaces exist^49^. Given we observed both dimers and higher-order oligomers, we cannot rule out that D_2_R interacts through additional interfaces depending on whether it exists in a dimeric or higher-order complex. Mutations at specific sites of the interface predicted herein increased the stability of D_2_R homomers, with the V96S/V97C double mutant exhibiting the greatest shift towards D_2_Rs in an oligomeric state in the absence of ligand (∼6% increase in all oligomer types compared to D_2_R WT), indicating an increase in constitutive oligomer stability. Super-resolution, single molecule analysis revealed there was little change in the proportion of WT and V96S/V97C D_2_Rs in an oligomeric state after cells were stimulated with quinpirole. Interestingly, the proportion of V96S receptors in an oligomeric complex at the cell surface were reduced when stimulated with quinpirole, which could reflect that di/oligomers of this D_2_R are internalized, consistent with the enhanced ligand-induced internalization and oligomerization favouring βarr2 coupling.

Ligand bias, whereby ligands can promote one pathway over another, are of interest both therapeutically and for understanding the molecular basis of pathway preferences for different GPCRs, in different conformational states^58,59^. In this study, receptor mutants, which showed increased di/oligomerization also showed increased ligand-induced (V96C and V96S) or constitutive (V96S/V97C) associations with βarr2 (Summarised in Fig. 6c-e). It is important to note that while the BRET assay results presented here suggest a steady state recruitment of βarr2, it cannot be ruled out that this represents multiple GPCR-βarr2 associations and dissociations, perhaps with distinct D_2_R-βarr2 conformations^60^.

There are many factors that can impact the signaling pathway preference of a GPCR, including membrane lipid composition and localisation of receptors^35,61–64^. Interestingly, these factors have also been shown to influence oligomerization of GPCRs^65^. While changes in the organisation of D_2_R complexes at the cell surface detected in PD-PALM experiments may be small, even slight changes in complex formation can have dramatic effects on receptor signaling in cells^37,38^. By enhancing specific receptor conformations such as a propensity to oligomerise and associate with βarr2, the role of such conformations on receptor activity may be amplified. These changes are in line with prior studies that have employed PD-PALM to assess GPCR di/oligomerization, where even small changes in protomer composition within a single oligomeric complex can direct how these receptors modulate signaling^37,38^.

There is some debate about the role of βarrs in D_2_R-mediated ERK activation. It is likely that mechanisms are both cell-type specific and D_2_R-isoform specific, with D_2_R long and short differing in the way they mediate MAPK activation^66,67^. Interestingly, the apparent bias towards βarr2 recruitment in D_2_R mutants results in enhanced internalization a βarr2-dependent mechanism for downstream mediation of the MAPK pathway. It is possible that increased dimerization of these mutants makes them more likely to internalize as has been demonstrated from CXCR4 oligomers^19^. Taken together, the most striking changes in signaling under basal conditions are observed with the D_2_R V96S/V97C double mutant which shows a strong preference for βarr2, required for downstream signaling. Here we have shown that different D_2_R mutants exhibit distinct βarr2 signaling profiles, supporting the broad spectrum of GPCR-βarr conformations that are being uncovered^68^.

The impact of oligomerization on biased signaling has focused on heteromers rather than homomers, including the A_2A_R-D_2_R heteromer^69,70^. Previous studies have reported that oligomerization can either increase or decrease G-protein activation, suggesting there is no consensus for all GPCRs in one class, but such behaviour is receptor specific^19,21,71,72^. The role of GPCR oligomerization on βarr recruitment has been less well studied. Oligomerization of the PAFR has been shown to cause a bias away from βarr recruitment, while our study demonstrates an opposing role for the D_2_R^21^. It has been suggested that allosteric interactions between receptor protomers upon receptor activation of the A_2A_R-D_2_R heteromer, causes the D_2_R protomer to favour βarr2 recruitment^70^. It is possible that a similar phenomenon is occurring in the D_2_R mutants explored in this study that exhibit increased βarr2 recruitment and associated signaling, also exhibit enhanced homomer stability.

Dimerization has been suggested to significantly increase the surface area available for arrestins to bind, which may provide a rationale for the increased oligomerization, which we observe in receptor mutants that also show a bias towards arrestin-mediated signaling pathways^11,73^. Consistently, the predicted D_2_R dimer is able to simultaneously accommodate two βarr2 molecules. It is possible that the V96S/V97C double mutant and, to some extent, the V96S D_2_R mutant, are showing a more ‘active-like’ conformation in the absence of agonists, which causes them to associate more readily with βarr2, causing increased internalization and a dependence upon βarr2 for MAPK signaling. Previous studies have found that βarrs are involved in D_2_R-mediated ERK phosphorylation in the A_2A_R-D_2_R heteromer, but not when D_2_R is expressed alone^74^. It is also possible that for these D_2_R mutants, increased homo-oligomerization induces increased dependence upon βarr2 for signaling pathways. This is supported by docking simulations between D_2_R homodimers and βarr2, which showed a 2:2 receptor: βarr2 stoichiometry while showing a 2:1 receptor: Gi stoichiometry (Fig. 6a and b and Supplementary Fig. 13). This suggests that D_2_R dimers, and therefore mutations which stabilize homo-di/oligomers, favour βarr2 coupling over Gi coupling. These docking simulations may provide an insight into the structural basis behind the observed increase in oligomerization in D_2_R mutants that also exhibit increased βarr2 associations.

In conclusion, the role of D_2_R oligomers in disease makes these complexes attractive therapeutic targets^30,75–77^. The lack of high-resolution structures of the D_2_R homodimer has hindered our ability to exploit oligomers for therapeutic benefits. Here we suggest models for the D_2_R homodimer and experimental validation confirms a key role for H1 and H2 at the predicted D_2_R homodimer interface to increase dimer stability, and a bias towards βarr2. These mutant constructs could form the basis for structural studies capturing D_2_R homomers and specific D_2_R-βarr complexes. More broadly, this study provides a framework for engineering the stabilization of class A GPCR dimers and may suggest that D_2_R oligomers favour an ‘active-like’ conformation that promotes βarr recruitment.

## METHODS

### Molecular modelling

Prediction of likely architectures of D_2_R homodimers relied on a computational approach developed for quaternary structure predictions of transmembrane α-helical proteins, including a number of GPCRs, i.e. FiPD-based approach^78–80^. It consists of rigid-body docking using both a version of the ZDOCK program devoid of desolvation as a component of the docking score (v2.1)^81^, followed by employment of a membrane topology filter by the FiPD software to minimize false positives. A rotational sampling interval of 6**°** was set (i.e., dense sampling) and the best 4000 solutions were retained and ranked according to the ZDOCK score. Such solutions were then filtered according to the “membrane topology” filter, which discards all those solutions that violate the membrane topology requirements. The membrane topology filter, indeed, discards all the solutions characterized by a deviation angle from the original z-axis, i.e. tilt angle, and a displacement of the geometrical center along the z-axis, i.e. z-offset, above defined threshold values, which were 0.4 radians and 6.0 Å respectively. The filtered solutions underwent cluster analysis by using a 3-Å Cα-atom Root Mean Square Deviation (RMSD) followed by visual inspection of the cluster centers for selecting the final solutions on the basis of membrane topology indices and docking score. Symmetrical D_2_R homodimers holding both protomers either inactive or active and asymmetric inactive-active dimers were predicted. The following structures were probed as sources of D_2_R protomers: a) inactive-state structures: 6CM4, 6LUQ, and 7DFP (i.e. D_2_R in complex with risperidone, haloperidol, and spiperone, respectively), and b) active-state structures: 7JVR and 8IRS (i.e. the ternary complexes between heterotrimeric Gi and D_2_R in complex with bromocriptine and ritigotine, respectively). The best results in terms of reliable docking solutions (according to the membrane topology indices) were achieved, for the active-state dimers, by using D_2_R from 8IRS as a protomer, and, for the inactive-state dimers, by-using D_2_R from 6CM4 completed with the missing IL2 as a protomer. The employment of an active-state structural model of D_2_R was also instrumental in predicting supramolecular complexes between D_2_R dimers and either βarr1 or βarr2. The latter predictions were carried out by ZDOCK v3.0.2^82^ coupled with a FiPD-based filtering approach based on a distance cutoff between the DRY arginine (R132) of D_2_R and L/V71 in the finger loops of βarr1 or βarr2, respectively, followed by cluster analysis. In those docking simulations, the receptor was used as a fixed target while βarr1 or βarr2 was used as a probe. The structural model of receptor-bound βarr2 were achieved by comparative modelling (by Modeller^83^), using the cyoEM structures of βarr1 bound either to the β_1_-AR (PDB: 6TKO) or to the 5HT_2B_ receptor (PDB: 7SRS) as templates. Remarkably, the two βarr1 templates differed in the conformation of several loops including the finger loop, an essential recognition point for receptor binding. Such comparative modelling was necessary because none of the available βarr2 structures, co-crystallized with peptides from the C-tails of a number of GPCRs, represent real receptor-bound forms as far as conformation and arrangements of the loops in the central crest are concerned. Accordingly, the employment of those βarr2 structures in docking simulations with the D_2_R did not produce any reliable solution. The employment of two different structural models of βarr2 served to overcome in part the lack of cryoEM complexes of βarr2 with GPCRs and to take into account possible structural variability in the receptor-bound states of the same arrestin. For βarr1, the most complete versions of the cryoEM structures 6TKO and 7SRS were employed. Only the results of βarr2-D_2_R docking are shown here. Evaluation of the native-like feature of a docking solution was based on structural comparisons with the cryoEM receptor-βarr1 complexes (6TKO and 7SRS).

### Cell Culture and transfection

HEK293 cells (human embryonic kidney cells) and βarr1/2 knockout cells were maintained as a monolayer in T-75 cm2 flasks (Sigma). βarr1/2 knockout HEK293 cells were made by using CRISPR-Cas9 (kindly provided by Asuka Inoue, Univ. Tohoku). Cells were cultured in Dulbecco’s Modified Eagle’s Medium (DMEM) media (Gibco) containing 4.5 g/L glucose, 1% (v/v) L-glutamine and supplemented with 10% (v/v) FBS (Gibco), 1% (v/v) streptomycin/penicillin (Sigma) in a humidified atmosphere of 5% CO_2_ at 37°C. Cells were usually passaged every 3-4 days in a 1:10 ratio using 0.05% (v/v) trypsin (Gibco, UK) in phosphate buffered saline (PBS). HEK293 cells were seeded and cultured in 6 well plates 1 day before transfection in DMEM with 10% (v/v) FBS (Gibco), 1% (v/v) streptomycin/penicillin (Sigma). Cells were transfected at 80-90% confluency with Lipofectamine 2000 reagent (Invitrogen, UK).

### Plasmid DNA constructs

βarr1-YFP and βarr2-YFP plasmids were obtained from Frederic Jean-Alphonse (CNRS, Nouzilly). FLAG-D_2_R-Venus and Rluc8 plasmids were obtained from Jonathan Javitch. The FLAG-D_2_R-Rluc8 plasmid was used to obtain the D_2_R Cys mutant constructs using the Agilent Quikchange II site-directed Mutagenesis Kit according to the manufacturer’s instructions. Mutant D_2_R constructs were introduced into D_2_R-Venus plasmids via the In-Fusion Snap Assembly cloning kit (Takara) according to manufactures instructions and inserted into FLAG-D_2_R-Venus DNA using BamHI and Kpn1 restriction sites. See Supplementary table 2 for a list of plasmids used in this study.

### BRET saturation assays

Cells were transfected with constant amounts of D_2_R-Rluc8 plasmid DNA and increasing amounts of D_2_R-Venus (0-3 μg in 6 well plates) plasmid and assayed 48 hours post-transfection. Cells in suspension were plated in white 96 well plates in triplicate. Coelenterazine-h (Promega) was used at a final concentration of 5 μM. BRET emissions using the short-wavelength filter at 475 nm and long wavelength filter 535 nm were measured in 10 repeat cycles using a LUMIstarOPTIMA plate reader (BMG Labtech). Fluorescence of Venus-tagged receptor was measured in duplicate after excitation at 485 nm and emission at 535 nm.

For each condition, net BRET values were calculated (535 nm value/ 475 nm value) over 10 cycles, where BRET ratios for the negative control donor only condition were subtracted. Rluc8 values (475 nm emission value) were averaged over the 10 cycles and again for the duplicate repeats. Net Venus values were calculated by subtracting Venus measurements of donor only conditions. Net BRET values were plotted against Net Venus/Rluc8 using GraphPad Prism 10 and fitted with a non-linear binding curve assuming one site-specific binding.

### β-arr recruitment BRET assays

HEK293 cells were transfected with D_2_R-Rluc8 plasmids and βarr1/2-YFP constructs. 48 hours post transfection cells were resuspended in PBS. For BRET time course assays, Venus expression (excitation 485 and emission 535 nm) was measured. For basal BRET emissions, a separate set of duplicate cell suspensions were measured at 475 nm excitation and 535 nm emission wavelengths after addition of 5 μM coelenterazine-h (Promega). PBS or 10 μM quinpirole was added to cell suspensions and BRET emissions were immediately measured again. Raw BRET ratios were calculated (535 nm value/ 475 nm value). To calculate ligand induced BRET, basal BRET ratios were subtracted from BRET ratios after agonist addition. For dose response BRET experiments, increasing concentrations of quinpirole agonist was added to sets of duplicate cell suspensions after basal BRET emissions had been measured. After ligand induced BRET values had been calculated, data was normalised by subtracting the control with only PBS added.

Analysis was based on βarr recruitment waveform methods as previously described^84^. The BRET emissions were measured every 3-5 seconds for a total of 5 minutes.

Ligand-induced BRET ratios were calculated and plotted against time and recruitment of βarr was observed to be stable recruitment, where the waveform rises to a steady state. The waveforms were fitted to non-linear one-phase association curve in GraphPad Prism 10 to allow quantification of kinetics. The rate constant s-1 was quantified and can be represented as the halftime, in seconds, (computed as ln(2)/k). The steady-state level of recruitment is represented as the plateau of the waveform, which also indicates the affinity of the receptor for βarr. The initial rate of recruitment was also calculated as k*span (maximum BRET ratio).

### Flow cytometry

Cells were live labelled with M1 anti-FLAG antibody (1:1000 Sigma Aldrich, no.3040) 48 hours post transfection for 30 minutes at 37 °C. For the flow internalization assay, antibody incubation was carried out at 4°C for 1 hour for all cells, then one set of cells were transferred to the 37 °C incubator and allowed to recover for 1 hour. Agonist stimulation at 37°C was carried out 30 minutes in to the 1-hour recovery period. The other set of cells was kept at 4 °C and washed 3 times with cold DMEM media. Cells were resuspended in FACS buffer (PBS with Ca^2+^and Mg^2+^ and 2% FBS), centrifuged and the cells resuspended in either FACS buffer alone for untreated cells or with Alexa Fluor 647 antibody (1:2000; Invitrogen, A32728). Cell suspensions were incubated for 1 hour on ice and washed in FACS buffer three times. The cell suspensions were transferred to individual round bottomed polypropylene tubes and the immunofluorescence measured on a FACS Calibur Flow Cytometer (BD biosciences).

### cAMP assay

HEK293 were transfected with 1 μg of D_2_R-Rluc8 constructs. After 24 hours, approximately 40,000 cells/well were replated into a 96-well format in triplicate for each condition. After a further 24 hours, the cells washed with PBS and pre-treated with 0.5 mM IBMX in DMEM with 0.1% Bovine serum albumin (BSA) for 5 minutes at 37°C. Forskolin was diluted in 0.5mM IBMX in DMEM 0.1% BSA to give a final concentration of 3 μM. Cells were stimulated with ligands diluted in the forskolin solution and incubated at 37°C. After stimulation, cells were washed with ice cold PBS and lysed. The level of cAMP in lysates was measured using the cAMP Gs Dynamic 2 kit (Revvity).

### Western blotting

Cells were washed with PBS and lysed with ice cold RIPA lysis buffer (50 mM Tris pH7.4, 1% Triton X-100, 140 mM NaCl, 5 mM EDTA, 1 mM NaF, 1 mM PMSF, 1 mM sodium orthovandate and a protease inhibitor tablet (Roche)). Protein concentrations were determined using BSA standard curve and diluted in Laemmli sample buffer (Sigma) in the presence or absence of 5% 2-β-mercaptoethanol. Samples were separated via SDS-polyacrylamide gel and transferred to a nitrocellulose membrane. The membrane was blocked and incubated in primary antibody diluted overnight (See Supplementary Table 1). Membranes were incubated in HRP-linked secondary antibody and imaged using Immobilon Forte Western HRP reagent (Millipore) and X-ray film (Amersham) or Chemi-imager. The band intensities were analysed using Image J.

For pERK Western blotting, cells were serum starved for 16 hours before quinpirole stimulation and lysis of cells. All lysates were diluted to 20 μg of protein in Laemmli buffer in the presence of 5% 2-β-mercaptoethanol, boiled for 5 minutes at 95°C before loading. Quantification of protein intensity was done separately for each blot and where multiple blots were used the same D_2_R WT lysates were employed to allow direct comparison with mutant D_2_R on the same membrane.

### Confocal Microscopy

Transfected cells were replated onto coverslips in a 24 well format with approximately 75,000 cells/well and left for a further 24-48 hours. Cells were incubated in M1 anti-FLAG antibody (Sigma Aldrich, no.3040) diluted 1:500 with or without 10 μM quinpirole for 30 minutes at 37°C. To remove the cell surface antibody, cells were washed 3 times with 0.04 M EDTA in PBS-Ca^2+^ before fixation. Cells were then fixed with 4% PFA in PBS-Ca^2+^, permeabilised with 0.2% Triton-X in PBS-Ca^2+^ and blocked for 1 hour at RT in 2% FBS in PBS-Ca^2+^prior to incubation in Alexa Fluro-conjugated secondary antibody (1:000 dilution) (Thermofisher, A32728). Slides were mounted onto coverslips and imaged were obtained with a Leica Stellaris 8 microscope using a 64 x oil immersion objective and sequential excitation at wavelengths of 405, 488 and 647 nm.

### PD-PALM

Transfected cells were plated onto 35mm dishes (Mattek) with 14mm x 1.5mm glass coverslips. Anti-FLAG (Sigma Aldrich) primary antibodies were labelled with photo switchable dye CAGE 500 according to the manufacturer’s instructions to give a 1:1 antibody:dye molar ratio (Abberior). Live cells were incubated with FLAG-CAGE500 conjugated antibody, diluted in 10% FCS in PBS Ca2+, at 37 °C for 30 minutes. All steps from antibody addition to imaging were carried out in the dark to prevent activation on the conjugated antibodies. Cells were washed in PBS Ca2+ before fixation in 4% PFA and 0.2% Glutaraldehyde for 30 minutes (Sigma) and maintained in PBS Ca^2+^.

PALM images were acquired using the Zeiss Eyra PS1 microscope with 1.45 numerical aperture, 100x oil immersion objective in total internal reflection fluorescence (TIRF) mode. Photoactivation of CAGE 500 dye was achieved by simultaneous activation with 405 nm (UV light) and excitation and photobleaching with 488 nm laser. Lasers were switched on 30 minutes before imaging to allow stabilization. Images were captured over 20,000 frames with an exposure time of 100 ms using the ZEN software.

Location analysis of receptors was done using the QuickPALM plugin in Fiji, as previously described ^85^. Two 7×7 μm region of interests were analysed within the cell boundary using the QuickPALM plugin. The parameters used were typically a signal to noise ratio of 7 and a full-half width maximum of 5, although these were determined for each experiment. This analysis outputs XY coordinates of the localised particles with sub-pixel accuracy. Particles within 20 nm of each other were discounted to prevent overestimation of receptor complexes. The coordinates were inputted into a Java-based PD-interpreter application ^38,39^. to determine the number of associated receptor molecules. This app uses Getis-Franklin neighbourhood analysis with a search radius of 50 nm. The software discounts a receptor from further searches once it has been assigned as participating in a homomer, to prevent double counting. The output is a table containing both the number of associations and the number of receptor molecules in each complex.

### Quantification and statistical analysis

All statistical analysis was performed in GraphPad Prism 10. Specific statistical tests are noted in each figure legend. In most cases, two-tailed, unpaired t-tests was used when comparing two groups or one-way ANOVA with Dunnett’s multiple comparison test was used for multiple groups. For comparing two variables and multiple groups, two-way ANOVA followed by Šídák’s multiple comparisons test was used. In all tests, p<0.05 was considered statistically significant.

## DATA AVAILABILITY

Source data are provided with this paper. Any additional data that support the findings of this study are available from the corresponding author upon reasonable request.

## CODE AVAILABILITY

Custom codes used to analyse PD-PALM data are provided as an attached Zip file.

## Supporting information

PD-PALM analysis software

## Acknowledgements

We would like to acknowledge Drs Stephen Rothery and David Gaboriau at the Facility for Imaging of Light Microscopy at Imperial College London for their technical support for microscopy experiments. Dr. Asuka Inoue (Univ. Tohoku, Japan) for supplying the β-arrestin1/2 KO cell line. Frederic Jean-Alphonse (CNRS, Nouzilly) for β-arrestin-YFP plasmids, Jonathan Javitch (Univ. Columbia, USA) for WT D_2_R plasmids. We would like to acknowledge the use of Biorender.com for the production of schematic figures. This project was funded by Biotechnology and Biological Sciences Research Council (Grant BB/M011178/1).

## Author contributions

B.B and A.C.H conceived the study and with K.L.S designed the experiments. K.L.S performed the majority of wet laboratory experiments, analysis and figure preparation with contributions from Y.L for certain βarr BRET recruitment assays and molecular biology; A.J.M for certain cAMP assays; W.Y for some confocal microscopy imaging.

F.F performed all molecular modelling and analysis. K.L.S wrote the majority of the manuscript with A.C.H and B.B. F.F wrote the sections of the manuscript relating to molecular modelling, with inputs and review by K.L.S, B.B and A.C.H. All authors contributed to and approved the final manuscript.

## Notes

### Competing Interest Statement

The authors have declared no competing interest.

### Summary of Updates

In this revised manuscript we have included new data with a distinct D2R-selctive ligand, additional controls for BRET data, a bias plot and modified the presentation of data to aid interpretation and improve clarity of results.

