## Supplementary Information for "Stabilized D_2_R G-protein coupled receptor oligomers identify multi-state β-arrestin complexes"

**Inventory**

**Supplementary Table 1. Antibodies used in this study**

**Supplementary Table 2. Plasmids used in this study.**

**Supplementary Fig. 1. Western blot analysis of HEK 293 cell lysates expressing D2R under non-reducing conditions.**

**Supplementary Fig. 2.** **D_2_R homomer interactions via BRET and expression levels of D_2_R mutants in HEK 293 cells**

**Supplementary Fig. 3. BRET saturation assays of D2R homomers in HEK293 cells following stimulation with quinpirole.**

**Supplementary Fig. 4. Models of predicted D_2_R homodimers**

**Supplementary Fig. 5. PD-PALM imaging of cells exhibiting either low or high receptor density to quantify oligomer populations.**

**Supplementary Fig. 6. Comparison of quinpirole-induced Gai signaling in either untransfected orD_2_R WT expressing HEK293 cells.**

**Supplementary Fig. 7.  βarr2, but not βarr1, is recruited following agonist activation of D_2_R WT.**

**Supplementary Fig. 8. Agonist-induced kinetics of βarr2 recruitment to wildtype (WT) and mutant D_2_R.**

**Supplementary Fig. 9. Lower receptor expression does not induce increased**

**basal βarr2 recruitment or increase either constitutive or ligand-induced internalization**

**Supplementary Fig. 10. Quinpirole and UNC9994 induce distinct D_2_R mediated G-protein signalling and β-arr-2 recruitment profiles.**

**Supplementary Fig. 11. Plasma membrane localization of WT and mutant D_2_R assessed via confocal microscopy.**

**Supplementary Fig. 12. Quantification of western blots assessing quinpirole-induced ERK1/2 signaling of wildtype and mutant D_2_R.**

**Supplementary Fig. 13. Predicted model for D2R homodimer-Gαi complexes.**

**Supplementary Fig. 14. BRET saturation assays of D_2_R homomers in HEK293 and βarr1/2 knockout cells**

**Supplementary Table 1. Antibodies used in this study.**

| **Antibody name** | **Type** | **Company** | **Product number** | **Dilution** |
| --- | --- | --- | --- | --- |
| Anti-GAPDH | Primary | Sigma | MAB374 | 1:1000 |
| HRP-linked horse anti-mouse antibody | Secondary | Cell Signaling | 7076 | 1:2000 |
| M1 Anti- FLAG | Primary | Sigma | 3040 | 1:1000 |
| Alexa Fluor Plus 647 antibody | Secondary | Invitrogen | A32728 | 1:2000 |
| HRP-linked mouse anti-rabbit | Secondary | Santa- Cruz | sc-2357 | 1:2000 |
| Anti- Alpha-tubulin | Primary | Cell Signaling | 2125 | 1:1000 |
| Phospho-p44/42 MAPK (erk1/2) | Primary | Cell signaling | 9101 | 1:1000 |
| P44/42 MAPK (erk1/2) | Primary | Cell signaling | 9102 | 1:1000 |

**Supplementary Table 2. Plasmids used in this study.**

| **Plasmid** | **DNA Construct Source** |
| --- | --- |
| FLAG-D2R-Rluc8 (wildtype) | Johnathan Javitch, Columbia University |
| FLAG-D2R-Venus (wildtype) | Johnathan Javitch, Columbia University, Addgene #19966 |
| FLAG- D2R- Rluc8 (V96C) | Michele Poli, Imperial College London |
| FLAG-D2R-Rluc8 (V96S) | Aylin Hanyaloglu, Imperial College London |
| FLAG-D2R-Rluc8 (V96S/V97C) | Aylin Hanyaloglu, Imperial College London |
| FLAG-D2R-Rluc8 (V97C) | Aylin Hanyaloglu, Imperial College London |
| FLAG-D2R-Rluc8 (Y93C) | Aylin Hanyaloglu, Imperial College London |
| FLAG-D2R-Venus (V96S) | Aylin Hanyaloglu, Imperial College London |
| FLAG-D2R-Venus (V96C) | Aylin Hanyaloglu, Imperial College London |
| FLAG-D2R-Venus (V96S/V97C) | Aylin Hanyaloglu, Imperial College London |
| FLAG-D2R-Venus (Y93C) | Aylin Hanyaloglu, Imperial College London |
| β-arrestin2-YFP | Frederic Jean-Alphonse, CNRS, Nouzilly |
| β-arrestin1-YFP | Frederic Jean-Alphonse, CNRS, Nouzilly |

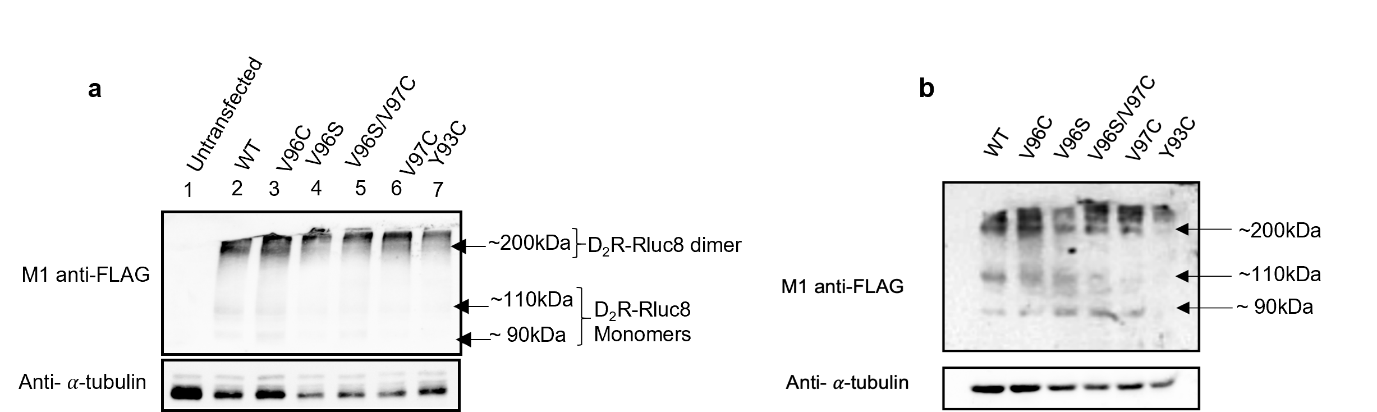

**Supplementary Fig. 1. Western blot analysis of HEK 293 cell lysates expressing D_2_R under non-reducing conditions.** Western blot of HEK293 cell lysates transfected with D_2_R-Rluc8 WT or mutant D_2_R-Rluc8 as indicated, carried out under non-reducing conditions. Membranes were probed with M1 anti-FLAG antibody to detect FLAG-tagged receptor and anti-α-tubulin was employed as a loading control. Two representative images from different biological replicates are shown and were imaged using a chemiluminescence imaging system (a) or using X-ray film (b).

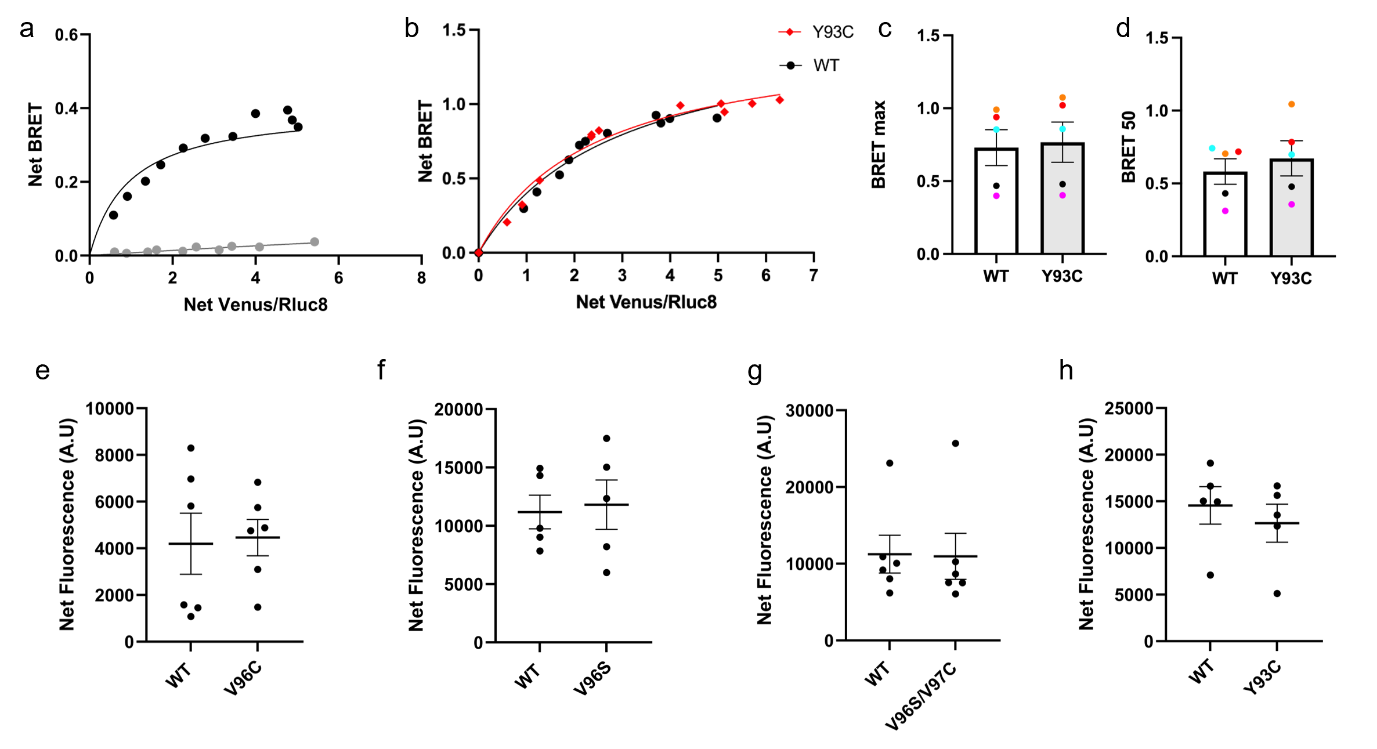

**Supplementary Fig. 2. D_2_R homomer interactions via BRET and expression levels of D_2_R mutants in HEK 293 cells.**  **(a)** Representative BRET saturation curve from HEK293 cells were transfected with constant amounts of D_2_R-Rluc8 WT with increasing amounts of D_2_R-Venus WT plasmid DNA (black). In grey, HEK293 cells transfected with increasing amounts of D_2_R-Venus and constant amounts of empty pcDNA3.1_Rluc8 vector plasmid was carried out to demonstrate specificity of saturation obtained in cells cotransfected with D_2_R-Rluc8 and D_2_R-Venus. **(b**) Representative saturation curves from HEK293 cells transfected with D_2_R WT (black) or D_2_R Y93C (red) plasmids. Curves used to calculate BRETmax (**c**) or BRET50 (**d**). N=5,+/- SEM.D_2_R-Venus WT or V96C (e), V96S (f), V96S/V97C (g) or Y93C (h) expressing HEK293 cells net fluorescence values. Cells were transfected with the same amount of D_2_R- Venus plasmid as used in Gi signaling, internalization and pERK assays. N=5 (V96S and Y93C) or 6 (V96C and V96S/V97C), +/- SEM.

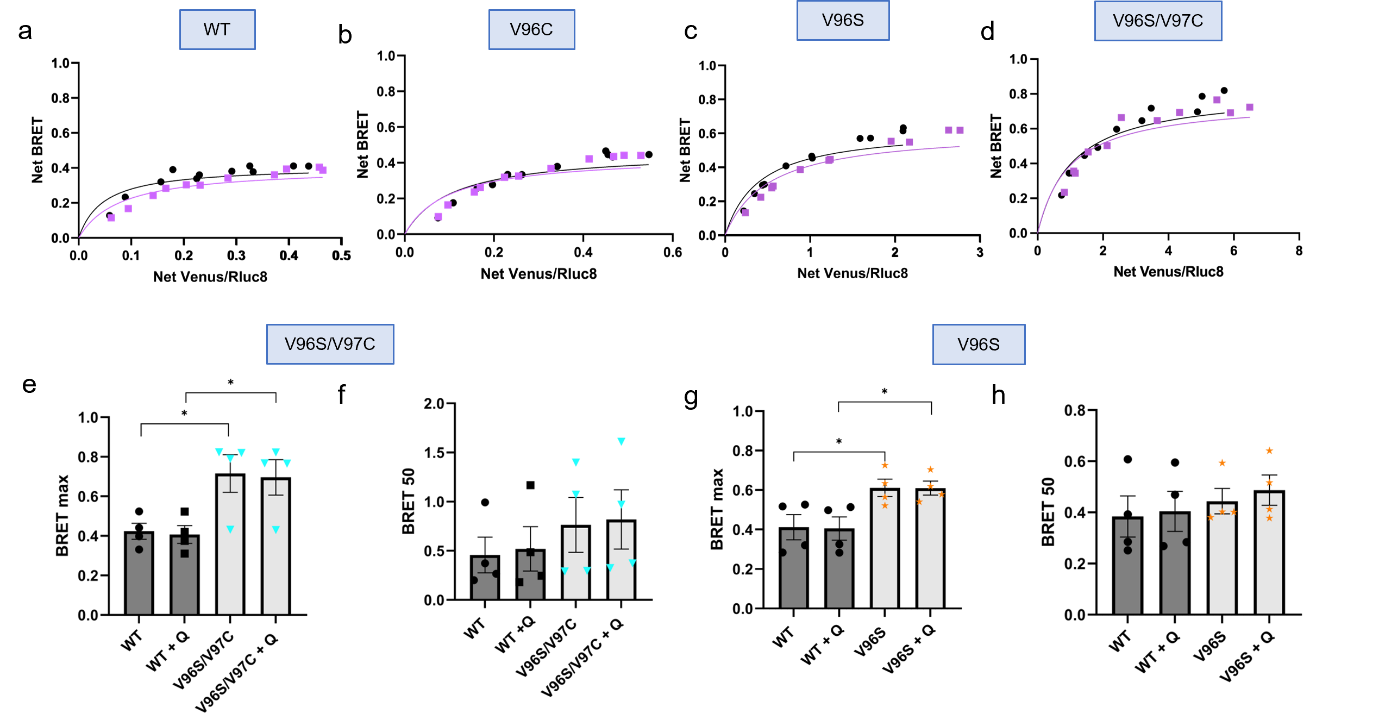

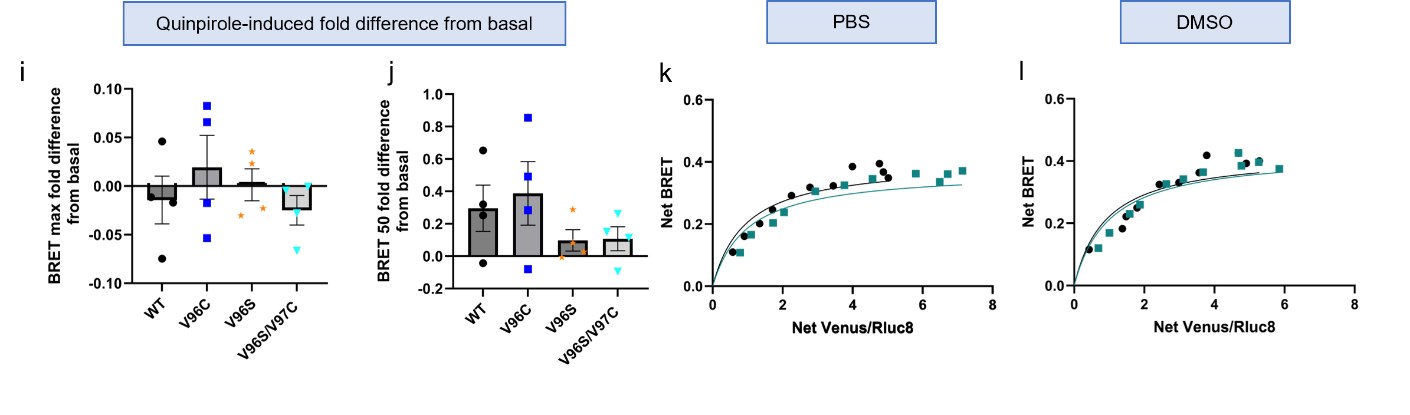
**Supplementary Fig. 3. BRET saturation assays of D_2_R homomers in HEK293 cells following stimulation with quinpirole.** HEK293 cells were transfected with constant amounts of D_2_R-Rluc8 WT with increasing amounts of D_2_R-Venus WT plasmid DNA (**a and e**) or mutant plasmids V96C (**b**) or V96S (**c**) or V96S/V97C (**d**). BRET signals were acquired before (black) and after addition of 10 μM D_2_R agonist quinpirole (purple). Saturation curves were used to quantify BRETmax (**e and** **g**) and BRET50 (**f and h**) and presented as a fold change from basal in I and j. N=4, +/- SEM. One-way ANOVA followed by Dunnett’s multiple comparison test used to compare basal and quinpirole treated and D_2_R WT and mutant expressing cells and across cell lines (**e**) p*=0.042 or 0.0245, (**g**) p*=0.0299 or 0.0281). (**k** and **l**) HEK293 cells were transfected with constant amounts of D_2_R-Rluc8 WT with increasing amounts of D_2_R-Venus WT plasmid DNA**.** Representative curves show the same transfected cell suspension as in Supplementary Fig.10 e and f before (black) and after (green) the addition of PBS (**k**) or DMSO (**l**) as controls.

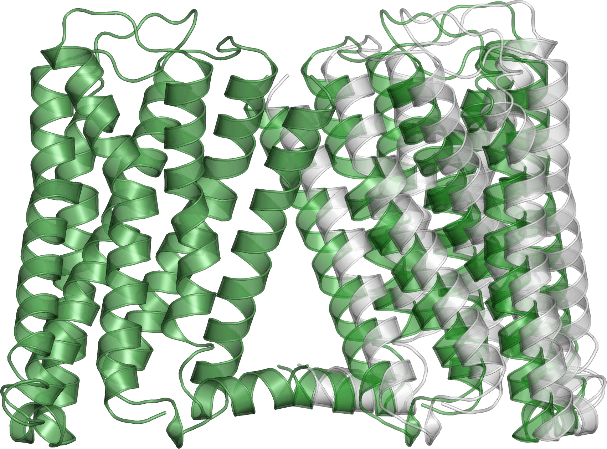

**Supplementary Fig. 4. Models of predicted D_2_R homodimers.** The cartoon representations of the superimposed D_2_R homodimers of WT (gray) and V96S/V97C mutant (forest-green) are shown.

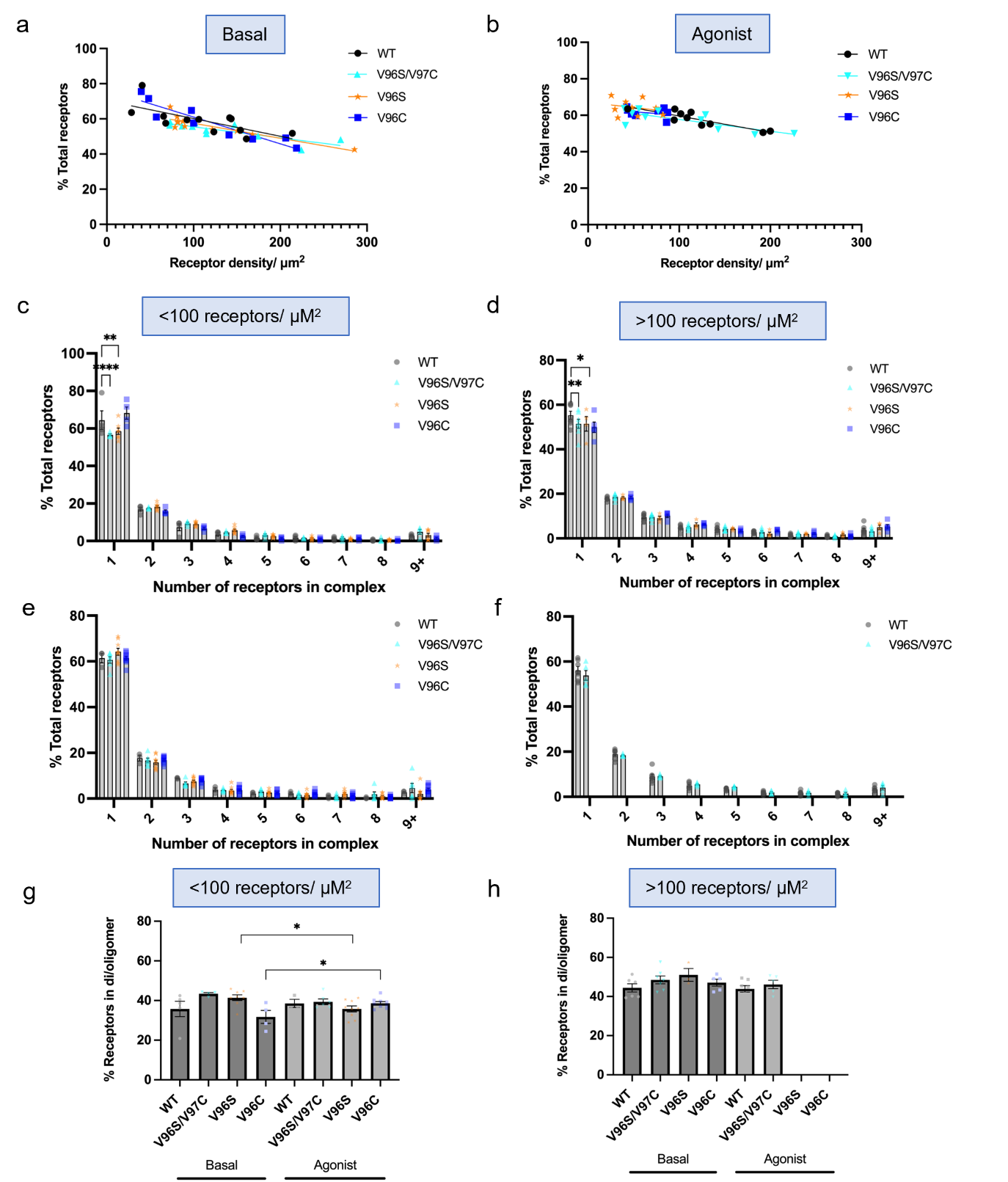

**Supplementary Fig. 5. PD-PALM imaging of cells exhibiting either low or high receptor density to quantify oligomer populations.** Correlation of D_2_R WT or mutant receptor density plotted against proportion of receptors that are not in an oligomeric state (monomers) without agonist stimulation (**a**) or after 5 minutes 10 μM quinpirole stimulation (**b**). Composition of D_2_R separated into number of receptors in a specific oligomeric complex (1-9 receptor molecules) and expressed as a percentage of total receptors in cells under basal conditions (**c and d**) or after 10 μM quinpirole stimulation **(e and f),** presented as cells with low (**c and e)** or high **(d and f)** receptor density. (**g and h**) Proportion of D_2_R WT or mutant receptors as monomers or in di/oligomeric complexes as a percentage of total number of receptors under basal conditions or after 5 minutes 10 μM quinpirole stimulation separated into cells with low or high receptor density. In c-f two-way ANOVA followed by Šídák's multiple comparisons test used to measure statistical differences between WT and mutant D_2_R in receptor complexes (p*=0.047 (d), p**=0.0011 (c) or 0.0013 (d), p****<0.0001). In g and h unpaired, two-tailed Student’s t test used to measure differences between WT and mutant D_2_R (p*= 0.0163 or 0.0164). N=3 individual experiments, 3-4 cells imaged for each independent repeat.

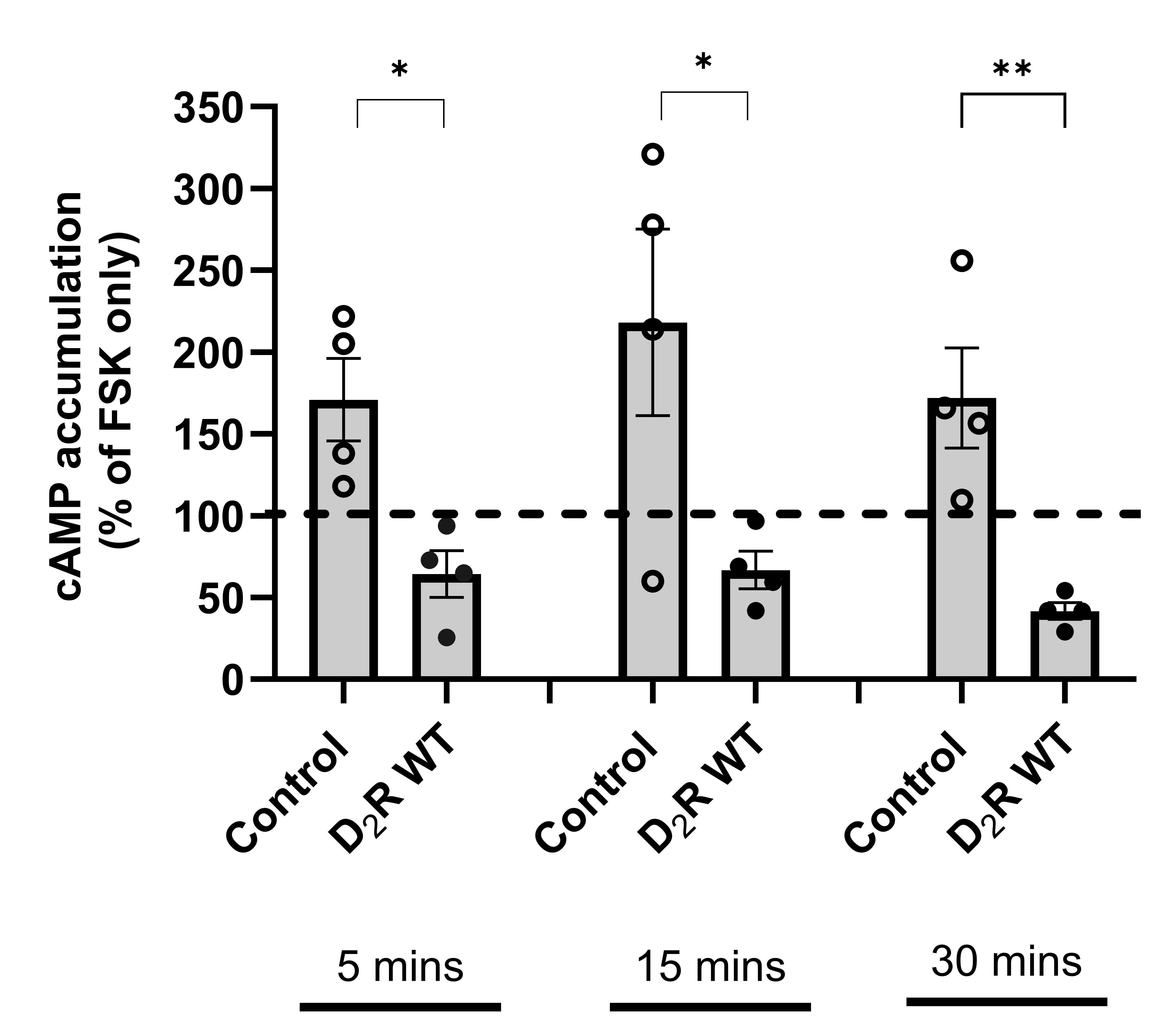

**Supplementary Fig. 6. Comparison of quinpirole-induced Gαi signaling in either untransfected or D_2_R WT expressing HEK293 cells.** cAMP accumulation of HEK293 cells transfected with D_2_R WT or untransfected (control) cells measured following 1 μM quinpirole stimulation in the presence of IMBX and forskolin for 5,15 or 30 minutes. Graphs represented as a percentage of cAMP levels in cells stimulated with forskolin only (100%), in the absence of agonist. Statistical analysis measuring differences to control untransfected cells within timepoints carried out with un-paired, two-tailed Student’s t test (p*=0.0104 or 0.0406, p**= 0.0057). N=4, error bars are +/- SEM.

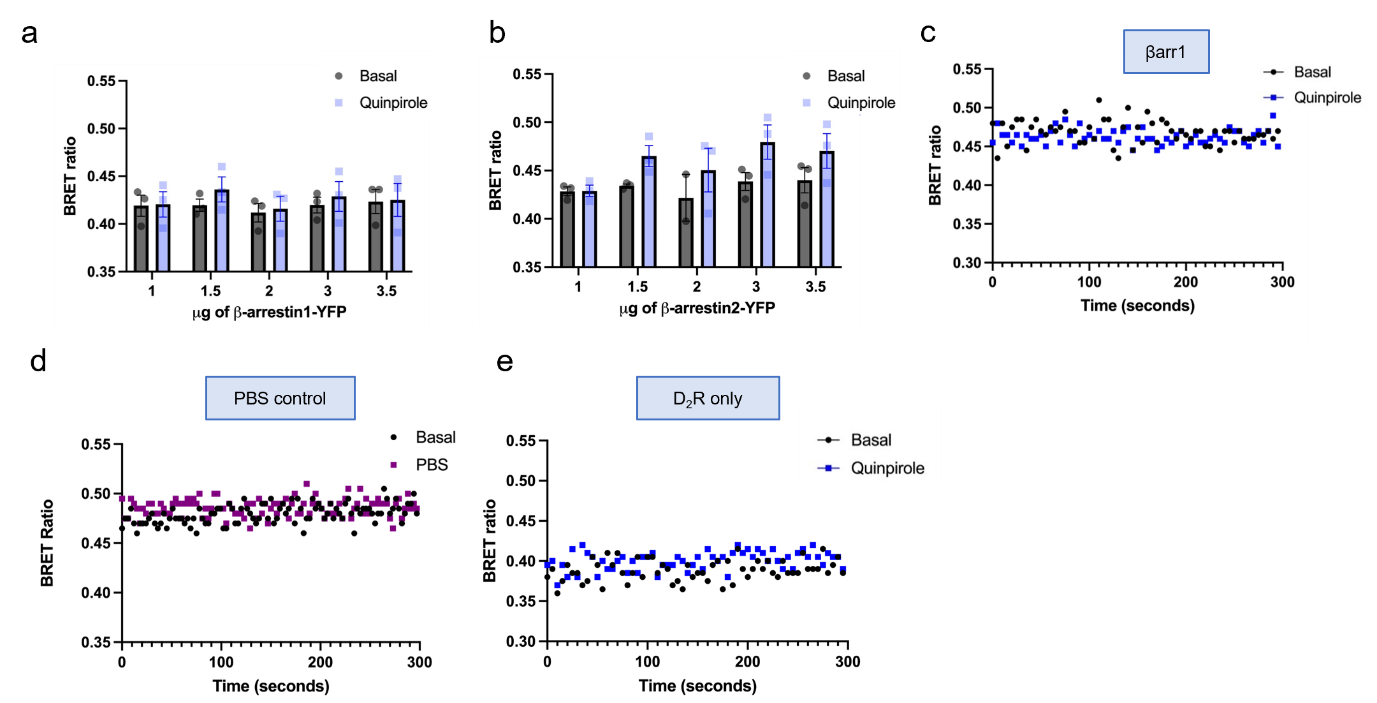

**Supplementary Fig. 7. βarr2, but not βarr1, is recruited following agonist activation of D_2_R. (a and b)** D_2_R-Rluc8 wildtype and increasing amounts of βarr1-YFP (**a**) or βarr2-YFP (**b**) were transfected into HEK293 cells and recruitment to the receptor was assessed before (black) and after (blue) quinpirole agonist addition by measuring BRET ratios. N=3, +/- SEM. **(c)** BRET measurements of cells transfected with D_2_R-Rluc8 WT and βarr2-YFP taken before and after addition of 10 μM quinpirole. **(d)** BRET measurements of HEK293 cells transfected with D_2_R-Rluc8 and βarr2-YFP with the addition of PBS (purple) instead of quinpirole after the first measurement, as a control. **(e)** BRET measurements of cells transfected with D_2_R-Rluc8 only and no βarr1/2-YFP before and after addition of 10 μM quinpirole. c, d and e show representative curves of at least 4 independent experiments.

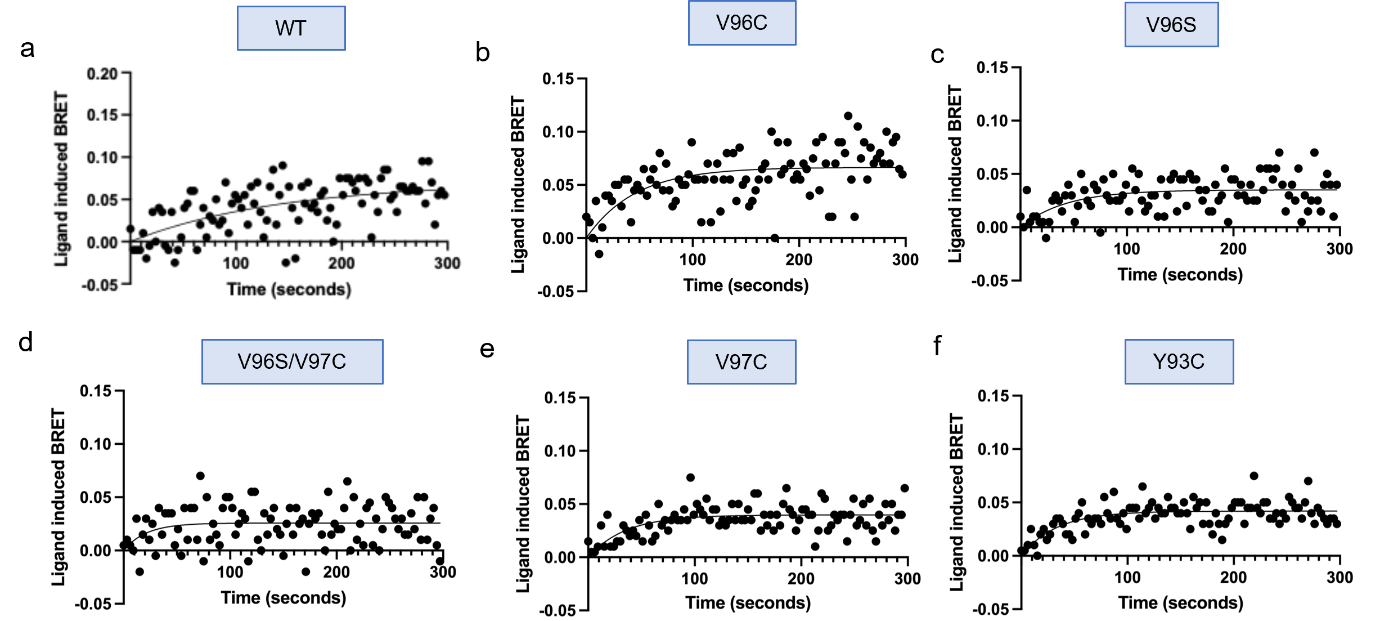

**Supplementary Fig. 8. Agonist-induced kinetics of βarr2 recruitment to wildtype (WT) and mutant D_2_R.** Representative kinetic profile of BRET signals in HEK 293 cells cotransfected with βarr2-YFP and D_2_R-Rluc8 WT (**a**), V96C (**b**), V96S (**c**), V96S/V97C (**d**), V97C (**e**), Y93C **(f**). Kinetics of curves produced from ligand-induced BRET values are quantified in Table 1. Ligand-induced BRET ratios calculated from BRET values in basal and quinpirole stimulated cells in D_2_R transfected cells as shown in Fig. 3. Graphs representative of 3-5 independent experiments.

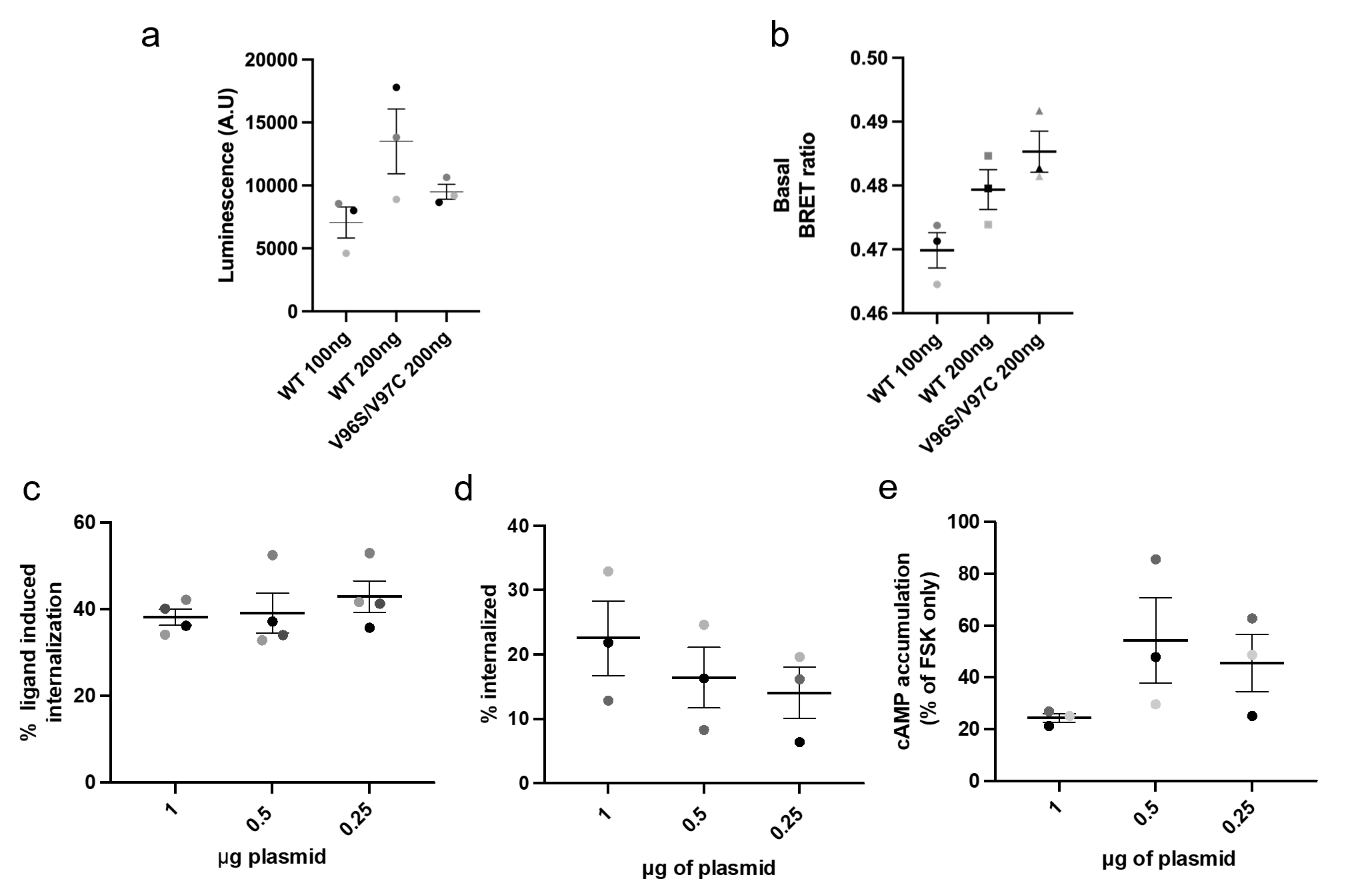

**Supplementary Fig. 9. Lower receptor expression does not induce increased**

**basal βarr2 recruitment or, increase either constitutive or ligand-induced internalization. (a)** D_2_R-Rluc8 luminescence values at 47 5nm wavelength after addition of coelenterazine-h to quantify receptor expression in HEK293 cells transfected with βarr2-YFP and 100 or 200 ng of D_2_R-Rluc8 WT plasmid or 200ng of D_2_R-Rluc8 WT plasmid (the same amount of D_2_R-Rluc8 plasmid DNA transfected in Fig. 3 and Supplementary Fig. 8). **(b**) Basal BRET values in D_2_R-Rluc8 and βarr2-YFP transfected cells, averaged over a timecourse of 5 minutes. N=3. Ligand induced internalization (**c**) and constitutive internalization (**d**) of WT D_2_R at lower expression levels (comparable to D_2_R V96S/V97C expression). Cell surface receptor expression assessed by flow cytometry and % internalization calculated as a decrease in cell surface receptor expression after 10 μM quinpirole stimulation ligand-induced internalization (**c**). For constitutive internalization (**d**), data presented as a percentage difference between cells incubated at 4°C (internalization prevented) and 37 °C (internalization can continue). N=3 or 4. (**e**) Forskolin-induced cAMP levels following 5 minute 100 nM quinpirole stimulation (taken as maximal response from dose response assays in Fig. 3a), presented as a percentage of cAMP levels in forskolin-only treated cells. Shaded dots correspond to individual biological replicates. N=3, +/- SEM for all experiments.

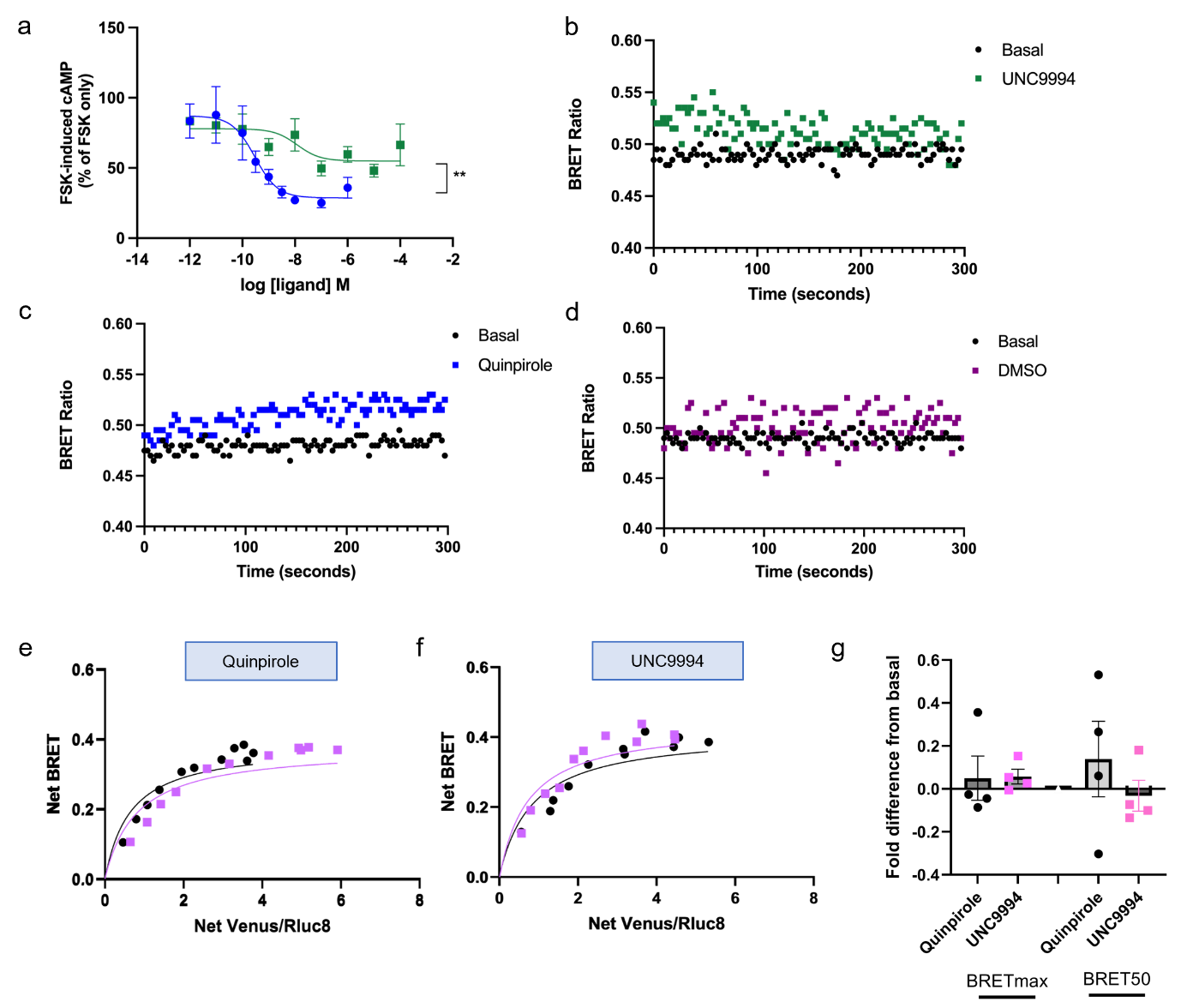

**Supplementary Fig. 10. Quinpirole and UNC9994 induce distinct D_2_R mediated G-protein signalling and β-arr-2 recruitment profiles. (a)** Dose response curves of forskolin-induced cAMP levels in the presence of IBMX in D_2_R-Rluc8 WT transfected cells following stimulation with UNC9994 (green) or Quinpirole (blue). N=3, +/- SEM, Statistical significance of differences in maximal Gi response assessed by un-paired Student’s t-test, **p= 0.0032. BRET assays showing kinetic profiles of β-arrestin-2 recruitment to D_2_R-Rluc8 WT before (black) or after 10 μM quinpirole (blue, **c**) or 10 mM UNC9994 (green, **b**) stimulation or with DMSO instead of ligand (purple, **d**) as a control. Presented as representative profile of 3 independent experiments. (**e-g**) HEK293 cells were transfected with constant amounts of D_2_R-Rluc8 WT with increasing amounts of D_2_R-Venus WT plasmid DNA**.** Representative saturation curves shown before ligand stimulation (black) or after stimulation with quinpirole (**e**) or UNC9994 biased agonist (**f**). (**g**) Saturation curves used to quantify BRETmax and BRET50 and changes following ligand stimulation presented as a fold difference from basal for each experiment. N=4, +/- SEM.

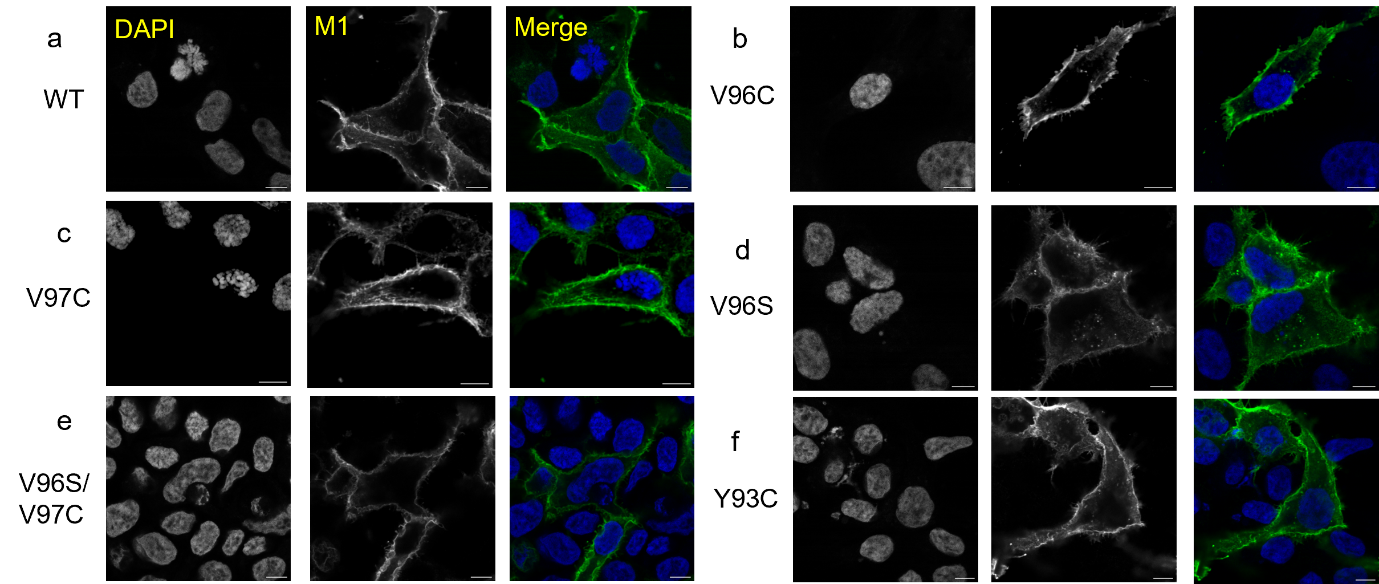

**Supplementary Fig. 11. Plasma membrane localization of WT and mutant D_2_R assessed via confocal microscopy** HEK293 cells transiently transfected with N-terminally FLAG tagged (**a**) D_2_R-Rluc8 WT, (**b**) V96C, (**c**) V97C, (**d**) V96S, (**e**) V96S/V97C, **(f**) Y93C treated with M1 anti-FLAG primary antibody and AlexaFluor647 secondary antibody, fixed and imaged using a confocal microscope to assess plasma membrane expression. Images representative of at least 3 independent experiments. Scale bar= 7 μm.

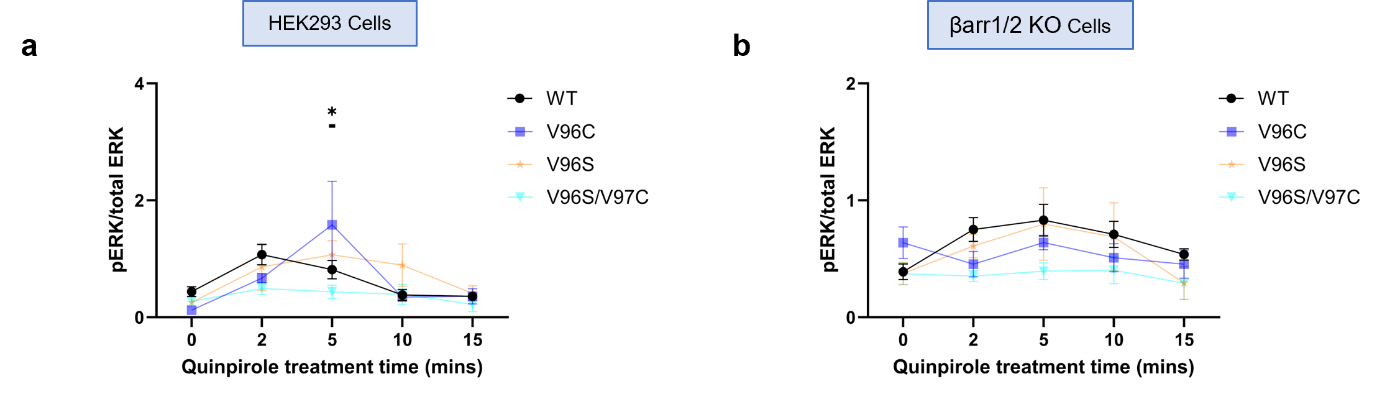

**Supplementary Fig. 12. Quantification of western blots assessing quinpirole-induced ERK1/2 signaling of wildtype and mutant D_2_R.** HEK293 cells (**a**) or HEK293 βarr 1/2 knockout (βarr 1/2 KO) cell lysates (**b**) transfected with D_2_R constructs and treated with 10 μM quinpirole for 2, 5, 10 or 15 minutes, were analysed by western blot and probed with phospho-ERK 1/2 and total ERK 1/2 antibodies. Represented as phospho-ERK1/2/total ERK protein levels. See Fig. 5 for representative western blots and data presented as a fold change from basal. N=8 for D_2_R WT expressing HEK293 and β-arr1/2 KO cells, N=6 for D_2_R V96C expressed in HEK293 cells, N=5 for D_2_R V96S and V96S/V97C expressed in HEK293 cells and D_2_R V96S expressed in β-arr1/2 KO cells, N=4 for D_2_R V96C and V96S/V97C expressed in β-arr1/2 KO cells. Error bars are +/- SEM. Two-way ANOVA followed by Šídák's multiple comparisons test used to measure statistical differences between WT and mutant D_2_R (in a WT vs V96C at 5 min timepoint,p*= 0.0356).

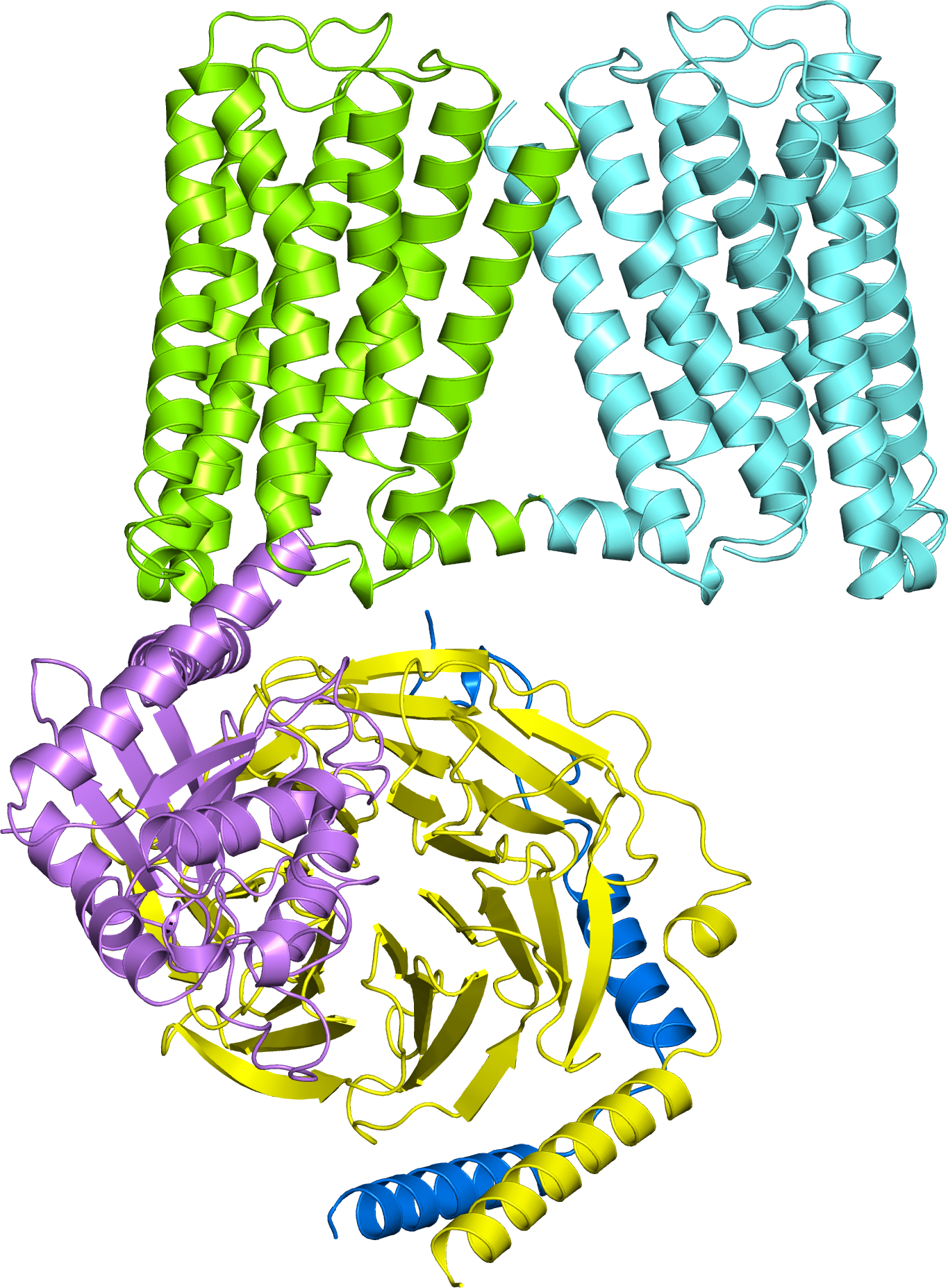

**Supplementary Fig. 13. Predicted model for D_2_R homodimer-Gαi complexes.** The cartoon representation of the predicted D_2_R WT homodimer (protomers lemon-green and aquamarine) and heterotrimeric Gi is shown, with a 2:1 receptor: Gi stoichiometry. The α, β, and γ subunits of Gi are shown in violet, yellow, and blue, respectively.

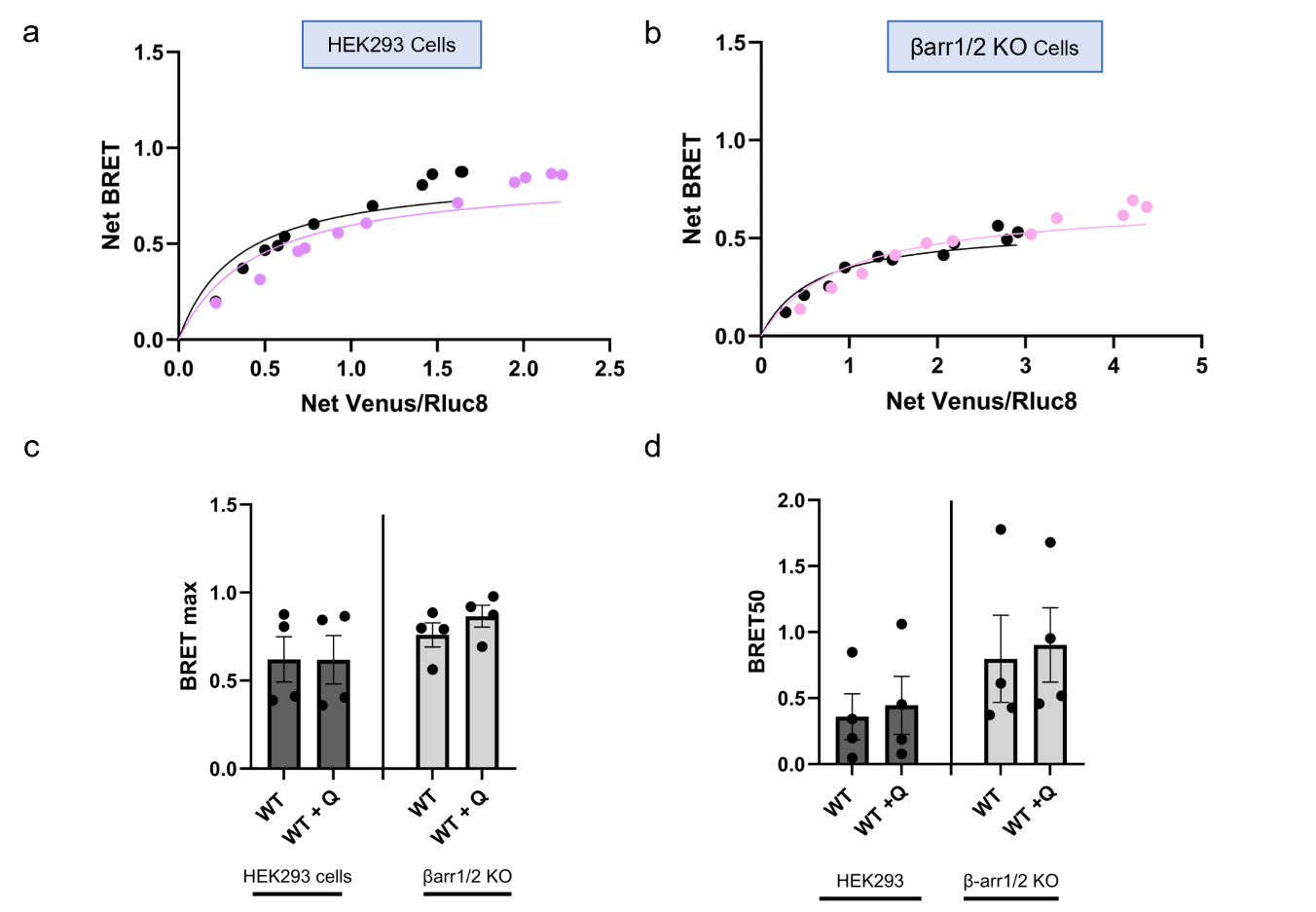

**Supplementary Fig. 14. BRET saturation assays of D_2_R homomers in HEK293 and βarr1/2 knockout cells.** HEK293 cells (**a**) and βarr1/2 knockout cells (**b**) were transfected with constant amounts of D_2_R-Rluc8 WT with increasing amounts of D_2_R-Venus WT plasmid DNA.BRET measurements were taken before (black) and after addition of 10 μM quinpirole (purple). Saturation curves were used to quantify BRETmax (**c**) and BRET50 (**d** N=4, +/- SEM. Unpaired, two-tailed Student’s t test used to measure differences between test used to compare between cell lines.
